# Dynamic rewiring of super-enhancers shapes condition-specific genome regulation

**DOI:** 10.64898/2026.11.12.766071

**Authors:** Sai Ananya Pavani A, Shirshanya Roy, Manjari Kiran

**Author notes:** Email addresses of authors: Sai Ananya, Shirshanya Roy, Manjari Kiran.

## Abstract

Enhancers and promoters communicate through three-dimensional chromatin contacts to control cell-type-specific gene expression. While chromosome conformation capture has enabled genome-wide mapping of these contacts, and graph-based analyses of the resulting interaction networks have shown that super enhancers occupy highly connected positions. However, how the network positions of super enhancers and typical enhancers changes as cells transition between states remains unclear. In this study, enhancer-promoter networks from high-resolution Hi-C data were constructed, and network-based approaches were applied to characterize the regulatory architecture and its remodeling during differentiation, using Hi-C and HiChIP datasets from three differentiation systems: PMA-induced K562 differentiation, cardiomyocyte differentiation of h9 cells, and differentiation of primary foreskin keratinocytes.

The networks are modular, with the largest clusters consisting of promoters and both enhancer types. Super-enhancer constituents and typical enhancers occupy distinct topological positions. During differentiation, network structure is mainly reshaped by enhancer turnover rather than by enhancer reorganization. Among the persistent enhancers, super-enhancer constituents rewire more extensively than typical enhancers. Functionally, remodeled super-enhancers connect to promoters controlling cell type’s identity-defining programs, whereas typical enhancers connect to promoters involved in general metabolic and stress-response loci. These results provide evidence that super-enhancers act as a sensitive, identity-focused layer of chromatin communication. The present study also provides quantitative, network-level evidence for how super-enhancers shape the architecture of chromatin communication.

## 4. Introduction

Transcription factors (TF) establish and control cell-type-specific gene regulation. Many TFs are expressed in a cell-type-specific manner and bind to gene regulatory elements (GRE) in the non-coding genome to control gene activity (Grosveld, Staalduinen, and Stadhouders 2021). Several classes of GRE have been identified, including enhancers, promoters, boundary elements, and silencers (Oudelaar and Higgs 2021), of which enhancers and promoters are most extensively studied. Enhancers are DNA elements that increase the transcription rate of a target gene, regardless of their orientation or distance from it (Shlyueva, Stampfel, and Stark 2014). Super enhancers (SEs) are large domains comprising multiple enhancers (referred to as Super enhancer constituents (SEc) hereafter) that act cooperatively. SEs are associated with high levels of enhancer activity, enrichment of enhancer-associated chromatin features (such as H3K27ac), a high occupancy of master regulators and coactivators (Whyte *et al*. 2013), and drive substantially higher target gene transcription compared to typical enhancers. SEs are reported to be associated with genes that define cell fate (Hnisz *et al*. 2013) and are also implicated in cancer, where they acquire genetic variants and regulate the expression of key oncogenic drivers (Thandapani 2019).

One widely accepted model by which enhancers exert their regulatory effect on the target gene is looping, in which the enhancer is brought into proximity with its target promoter. Sequence-specific TF bound to the enhancer, together with architectural proteins such as cohesin and the mediator complex, establishes and stabilizes the loops, allowing contact with the promoter and recruiting RNA polymerase II, thereby boosting target gene transcription (Panigrahi and O’Malley 2021). Advances in chromosome conformation capture techniques, particularly high-throughput chromosome conformation capture (Hi-C), allow such interactions to be mapped genome-wide, and derivative techniques such as in situ Hi-C (Rao *et al*. 2014), capture Hi-C (Schoenfelder *et al*. 2018), and HiChIP (Mumbach *et al*. 2016) have extended this capability. Hi-C data is summarized as a matrix of pairwise contact frequencies, effectively describing the three-dimensional genome architecture as a many-to-many interaction network.

This network representation invites analysis with the tools of graph theory. A network comprises a set of nodes or vertices connected by edges or links, enabling the interpretation of complex systems through their relationships. A biological system can be represented as a network, with nodes representing biomolecules (such as DNA or proteins) and edges representing interactions between them, which may be physical, functional, or chemical in nature (Zitnik *et al*. 2024).

The importance and connectivity of individual nodes can then be quantified through centrality measures: degree captures the number of direct connections and strength (weighted degree) their cumulative interaction weight; closeness measures the average distance to all other nodes; betweenness quantifies how often a node lies on shortest paths between others; harmonic centrality measures a node’s reachability to the rest of the network; and eigenvector centrality measures influence weighted by the importance of a node’s neighbors (Newman 2018). Together they help distinguish different nodes.

Several studies have represented the Hi-C matrix as a graph, with interacting regions as nodes and contacts as weighted edges. (Belyaeva *et al*. 2017, Liu and Ruan 2017, Malod-Dognin *et al*. 2020). Network analyses of chromatin interactions have also been used to study specific classes of regulatory elements. Thiobdeau et al. (2017), for example, showed that super enhancers are highly connected than other enhancers (referred to as typical enhancers (TE) throughout the manuscript hereafter). (Thiobdeau *et al*. 2017). These studies, however, mostly describe a single cellular state, it still remains unclear whether SEc and TE keep or change their network positions when cells change state, and whether such changes affect the gene programs they regulate.

Here, we apply established network analysis methods to study enhancer-promoter communication during differentiation. We integrated significant interactions from high-resolution Hi-C data with SEc, TE, and promoter annotations to construct the enhancer-promoter interaction networks in which edges are weighted by the loop interaction frequency. For each enhancer class, we calculated the standard graph-theoretic centrality measures and compared them across conditions, enabling tracking of network reorganization during cell state transitions. By contrasting these metrics and identifying the functions of promoters engaged by SEc and TE-anchored loops that change across conditions, we test, from a network perspective, the long-standing hypothesis that super-enhancers are key determinants of cell fate and act as master regulators during cell state transitions. The present work aims to provide quantitative, network-level evidence for how super-enhancers shape the architecture of chromatin communication.

## 5. Data Collection

### 5.1 Hi-C data

Processed Hi-C data for the K562 cell line (control/treated) were collected in .hic format from Bond et al. (GSE no: GSE214123) (Bond *et al*. 2023). K562 cells represent the cell culture model of chronic myelogenous leukemia. Upon treatment with PMA (phorbol 12-myristate 13-acetate), K562 cells acquire a macrophage-like morphology (Rehemtulla, Arndt, and Hart 1990), providing a model of cellular differentiation. Accordingly, three conditions: one untreated and two PMA-treated (6 hrs and 72 hrs post-treatment) were selected for the primary analysis and are summarized in Table 1.

**Table 1.**
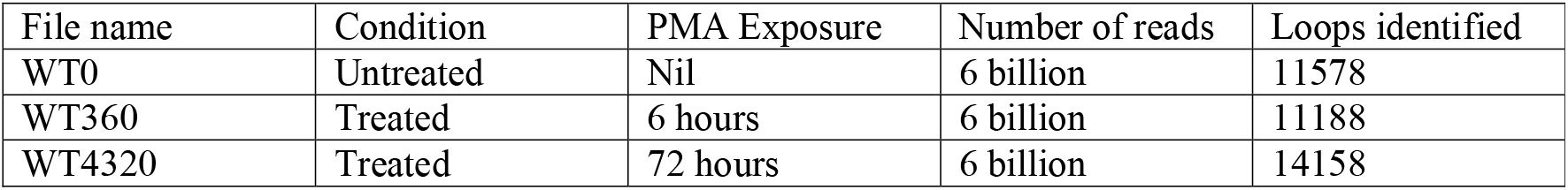
xxxxx

| File name | Condition | PMA Exposure | Number of reads | Loops identified |
|---|---|---|---|---|
| WT0 | Untreated | Nil | 6 billion | 11578 |
| WT360 | Treated | 6 hours | 6 billion | 11188 |
| WT4320 | Treated | 72 hours | 6 billion | 14158 |

To confirm our findings, the observed patterns were examined in independent Hi-C and HiChIP datasets from multiple sources. Hi-C data for the H9 differentiation series were obtained from the 4D Nucleome (4DN) Data Portal (Reiff *et al*. 2022; accession IDs: 4DNFII6AN691, 4DNFINEQY95T, 4DNFIJ8JKKWJ). Processed H3K27ac HiChIP loop calls for the K562 cell line and foreskin keratinocyte differentiation were obtained from the Loop Catalog (Reyna *et al*. 2025). Details of all datasets, including sample conditions and sources, are provided in Supplementary Table 1.

### 5.2 Promoter Coordinates

Regulatory Features and Regulatory Activity files were downloaded from Ensembl release 112 for K562 and foreskin, or release 114 for h9 (Cunningham *et al*. 2022, Harrison *et al*. 2024). The Regulatory Activity file indicates whether a given regulatory element is active, poised, repressed, or inactive in a specific cell type. IDs of active promoters specific to each cell line were extracted from the Regulatory Activity file, and their corresponding coordinates were taken by cross-referencing the Regulatory Features file.

### 5.3 Super-enhancer and typical enhancer coordinates

Coordinates of SEc and TE specific to all the cell types were downloaded from SEdb 3.0. (Song *et al*. 2026)

## 6. Hi-C Data Analysis

Significant chromatin loops were identified from Hi-C data using the command-line tool Significant Interaction Peak (SIP) caller (Rowley et al. 2020). SIP was performed at default parameters at 5 kb resolution. Interacting regions and normalized interaction frequency were then obtained from the output BED files for subsequent analysis.

For H3K27ac HiChIP data, loops were obtained from the Loop Catalog in BED format (Reyna *et al*. 2025), and interacting regions and interaction frequency columns were obtained for subsequent analyses.

## 7. Construction of networks using identified loops

Networks were created in R (v4.6.1) using loops identified from Hi-C or H3K27ac HiChIP datasets (R Core Team 2026). The following R packages were used: igraph (Csardi 2014), GenomicRanges (Lawrence *et al*. 2013), and InteractionSet (Lun, Perry, and Ing-Simmons 2016).

### 7.1 Annotation of loop anchors

To determine whether loop anchors overlap with promoters (P), SEc, or TEs, loop coordinates were converted into a GInteractions object (InteractionSet), and genomic coordinates of P, SEc, and TEs were converted into GRanges objects (GenomicRanges). Loop anchors were relaxed by 10 kb, and findOverlaps() was used to identify overlaps between each anchor and the three regulatory feature sets. Overlap results were compiled into a truth table with six logical columns (A1_P, A1_SE, A1_TE, A2_P, A2_SE, A2_TE), indicating whether anchor 1 (A1) or anchor 2 (A2) overlapped each regulatory element type. The IDs of all intersecting regulatory features were appended as separate columns for each anchor.

### 7.2 Classification of loop anchors and network generation

Each loop anchor was given a single regulatory element label based on the priority order: P> SEc> TE, following the priority used in the previous study (Lee, Wu, and Rhie 2022), and corresponding regulatory element ID(s) were assigned to each anchor. To confirm that the observed trends were not an artifact of the priority scheme used, the network was also constructed with a reversed priority order (TE > SEc > P), and network properties were calculated to assess whether annotation preference introduced any bias.

After both anchors of a loop were labeled, each chromatin loop was represented as a connection (edge) between regulatory elements, with edge weight defined as the interaction frequency of the loop. If an anchor overlapped several regulatory elements, all possible combinations between two anchors were calculated to ensure that no element-element interactions were omitted. This process produced an edge data frame, in which each row represented a unique element pair and its associated interaction frequency, and a corresponding node data frame containing element IDs and element type.

### 7.3 Network construction and properties

The edge and node data frames were used to construct the enhancer-promoter network using the igraph package. Connected components of the network were identified and were classified based on their constituent element types. For each cluster, network properties, including degree, strength, betweenness, closeness, harmonic centrality, and eigenvector centrality, were calculated for enhancer nodes (SEc and TE), with edge weights incorporated where applicable.

To track node partners of enhancers across conditions we calculated the jaccard similarity index between enhancer’s partner sets in the two conditions, for conciseness, we call this as the partner conservation index (PCI). for each enhancer *i* present in both conditions A and B:

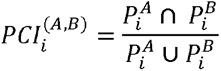

Where,

*P_i_*^A^ - set of interaction partners of enhancer i in condition A

*P_i_^B^* - set of interaction partners of enhancer i in condition B

### 7.4 Bipartite network projection

To determine whether the observed patterns are an intrinsic property of E-P wiring rather than a network projection artifact, a bipartite network was built from the Hi-C loops in the WT0 condition, retaining only edges connecting enhancers (SEc or TE) to promoters, along with their corresponding edge weights.

## 8. Functional enrichment of promoters targeted by dynamic SE-P and TE-P loops

To assess the effect of enhancer turnover during PMA-induced differentiation of K562 cells, two conditions were selected: 72 hours post-treatment (WT4320) and the untreated state (WT0). The loops identified in both conditions were merged into a union loop set. Loops that appeared only in WT4320 were designated as de novo gained, whereas those found only in WT0 were designated as de novo lost. For functional enrichment analysis, only enhancer-promoter interactions (SEc-P and TE-P) were considered from the union set. Promoter anchors participating in these interactions were extracted, and their genomic coordinates were compiled for each loop category.

Promoter functional enrichment was performed using rGREAT (Gu and Hübschmann 2023), with the MSigDB Hallmark gene set collection (Liberzon et al. 2015) as the reference database, against the genome. Multiple testing correction was applied to evaluate enrichment significance. The resulting enriched pathways were compared across SEc-P and TE-P categories to identify biological processes associated with SEc- and TE-mediated regulatory rewiring.

## 9. Results and discussion

### 9.1 Enhancer promoter networks are modular, with SEc and TE occupying distinct positions

Alt text: Four panels describing the enhancer-promoter network of untreated K562 cells (WT0), built from Hi-C loops. (a) Bar chart showing loop counts for each interaction category. Categories are labeled with colored squares indicating anchor types: promoter (brown), super-enhancer constituent (purple), or typical enhancer (teal). (b) Network diagram of the enhancer-promoter interaction network. Many separate modules are visible, rather than a single connected network. (c) Bar chart showing the number of connected components for each type: P-SEc, P-TE, and P-SEc-TE. (d) Box plots showing the cluster sizes for the three types described above.

Three experimental conditions were used to construct the network: untreated control (WT0) and two PMA-treated time points (WT360 - 6 hours after PMA treatment and WT4320 - 72 hours after PMA treatment). In the WT0 condition, 11578 loops were identified, of which approximately 12% of the loops were found to be involved in interactions between enhancers, promoters, and enhancer-promoter pairs (Figure 1a). This result is consistent with a previous study stating that the majority of observed interactions are between other regulatory elements, such as CTCF sites, and that loops constituting enhancer-promoter interactions constitute a small percentage of the total loops identified. (Lee, Wu, and Rhie 2022).

**Figure 1:**
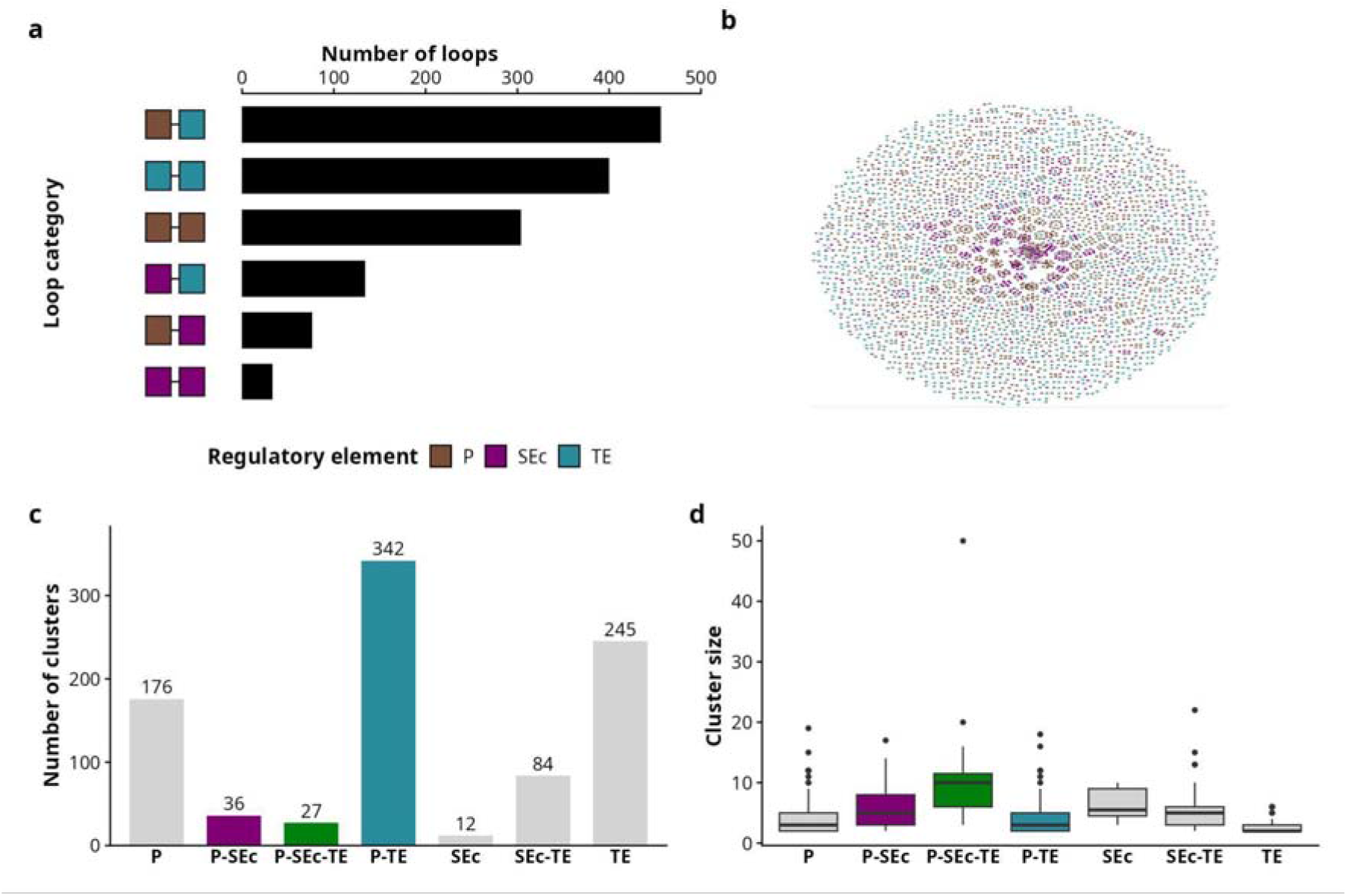
Representative diagram showing the loop and network characteristics: (a) Number of loops obtained in each interaction category; (b) Enhancer-promoter contact network of K562 cell line (untreated) generated from Hi-C data; (c) Number of clusters in each cluster category; (d) Size distribution of clusters within the network.

The baseline network comprised 1782 P, 645 SEc, and 1245 TE nodes (Table 2) and consisted of many connected components. The network had a high modularity score (modularity = 0.863, at a resolution of 1) (Figure 1b), suggesting that the enhancer-promoter interaction network possesses a modular regulatory architecture with enhancer-promoter interactions organized into distinct regulatory modules rather than being uniformly distributed throughout the network; such modular organization supports the highly specific nature of enhancer-promoter interactions and may also facilitate precise spatiotemporal regulation of gene expression by enabling coordinated regulatory activity within discrete components.

**Table 2.** xxxxx

| File name | Nodes | Edges | Typical Enhancers | Super Enhancers | Promoters | Largest Connected Component size | Number of Clusters |
|---|---|---|---|---|---|---|---|
| WT0 | 3672 | 4267 | 1245 | 645 | 1782 | 50 | 922 |
| WT360 | 3466 | 3557 | 1248 | 528 | 1690 | 46 | 903 |
| WT4320 | 3747 | 3843 | 1294 | 600 | 1853 | 33 | 929 |

Nodes comprising the connected components were identified, and clusters consisting of promoter and enhancer (SEc or TE) were considered to compare network properties of SEc and TE (Figure 1c). It was observed that the cluster size of the components comprising all three different kinds of nodes: TE, SEc, and P, on average, was larger than that of the components of other categories. (Figure 1d), and this trend was also retained in the bipartite network. (Supplementary Figure 1). The larger average size of P-SEc-TE clusters can be attributed to the fact that strongly regulated genes receive multiple signals from enhancers (Uyehara and Apostolou 2023), and cluster organization can help facilitate this.

To understand whether these observations were specific to the K562/PMA-treated K562 system or if they represented a broader feature of enhancer-promoter organization, additional enhancer-promoter networks were constructed from multiple independent datasets, including the H9 differentiation-series Hi-C datasets and H3K27ac HiChIP datasets from untreated K562 and foreskin keratinocyte cells. The observed trends remained consistent: networks constructed from the datasets were highly modular (Supplementary Table 1), and connected components comprising P, TE, and SEc exhibited larger cluster sizes on average than components belonging to other interaction categories (Supplementary Figure 2).

The centrality measures were calculated for SEc and TE nodes in the P-SEc, P-TE, and P-SEc-TE clusters, as these nodes represent the major regulatory interaction classes directly involving connections between enhancer elements and promoters. In untreated K562 cells, the trends observed between SEc and TE for degree, betweenness, and eigenvector centrality were consistent with earlier network analyses of super enhancers by Thibodeau *et al*. (2017), with SEc having higher degree and betweenness than TE, and TEs having higher eigenvector centrality than SEc. In addition to these metrics, strength, closeness, and harmonic centrality were also computed. Overall, SEc tend to be well-connected, have stronger interaction strength, and show higher harmonic centrality within the network compared to TE, suggesting that they act as local regulatory hubs of the E-P network that occupy distinct network positions relative to TE. (Supplementary Figure 3a).

Among networks constructed from other cell lines, SEc exhibited systematically higher local connectivity than TE. Degree, strength, and harmonic centrality were greater for SEs across all examined datasets and contexts. Global centrality measures favored TEs; eigenvector and closeness centralities were higher in TEs across most contexts and cell lines/datasets. (Supplementary Figure 3)

Among both classes of enhancer nodes, SEc are enriched for local connectivity metrics but depleted for global connectivity metrics such as eigenvector centrality and closeness, and show no preference for betweenness. This is consistent with the model in which SEs function as interconnected, local regulatory hubs.

Because E-P regulation operates predominantly as distinct modules, the local connectivity metrics on which SEc consistently dominate (degree, strength, and harmonic centrality) are the more biologically informative measures of regulatory influence, whereas global metrics favoring TE (eigenvector and closeness) reflect their embedding within the sparse network rather than direct regulatory control. The elevated eigenvector centrality of TE itself might reflect its attachment to high-degree SEc hubs, making this metric a downstream signature of SEc connectivity. Together, these topological features position SEc as the principal regulatory drivers of the network, with TE acting as the peripheral connectors.

The trends observed in the non-bipartite network were also largely seen in the enhancer-promoter bipartite network (Supplementary Figure 4), suggesting that the central role of SEc is an intrinsic property of E-P wiring rather than a projection artifact.

Key network trends remain largely preserved even after reversing the annotation hierarchy (Supplementary Figure 5), showing that the observed properties are not artifacts of the hierarchy used for node assignment and that these trends are robust to the annotation scheme. One particular trend is that even when TEs are given annotation priority, SEc-associated nodes still retain elevated degree/strength/harmonic centrality, which argues against the possibility that SEc centrality is simply caused by preferential annotation assignment. For HiChIP datasets as well, annotation priority does not affect network trends.

### 9.2 Enhancer turnover shapes network remodeling during differentiation

Alt text: Six box plot panels comparing various network centrality measures for super-enhancer constituents and typical enhancers across PMA treatment conditions in K562 cells. The panels show (a) degree, (b) strength, (c) betweenness, (d) closeness, (e) harmonic centrality, and (f) eigenvector centrality. In each panel, SEc and TE are separated by condition, with shading from dark to light for initial to later time points. Significance is indicated by asterisks above each enhancer type.

Since super-enhancers are known to play a role in cell fate determination, the differentiation could be accompanied by changes in the enhancer-promoter network. Therefore, the changes in network properties for SEc and TE across conditions were assessed. It was found that both SEc and TE showed changes in network properties across conditions (Figure 2a-f; reproduced in H9 and foreskin keratinocytes, Supplementary Figure 6).

**Figure 2:**
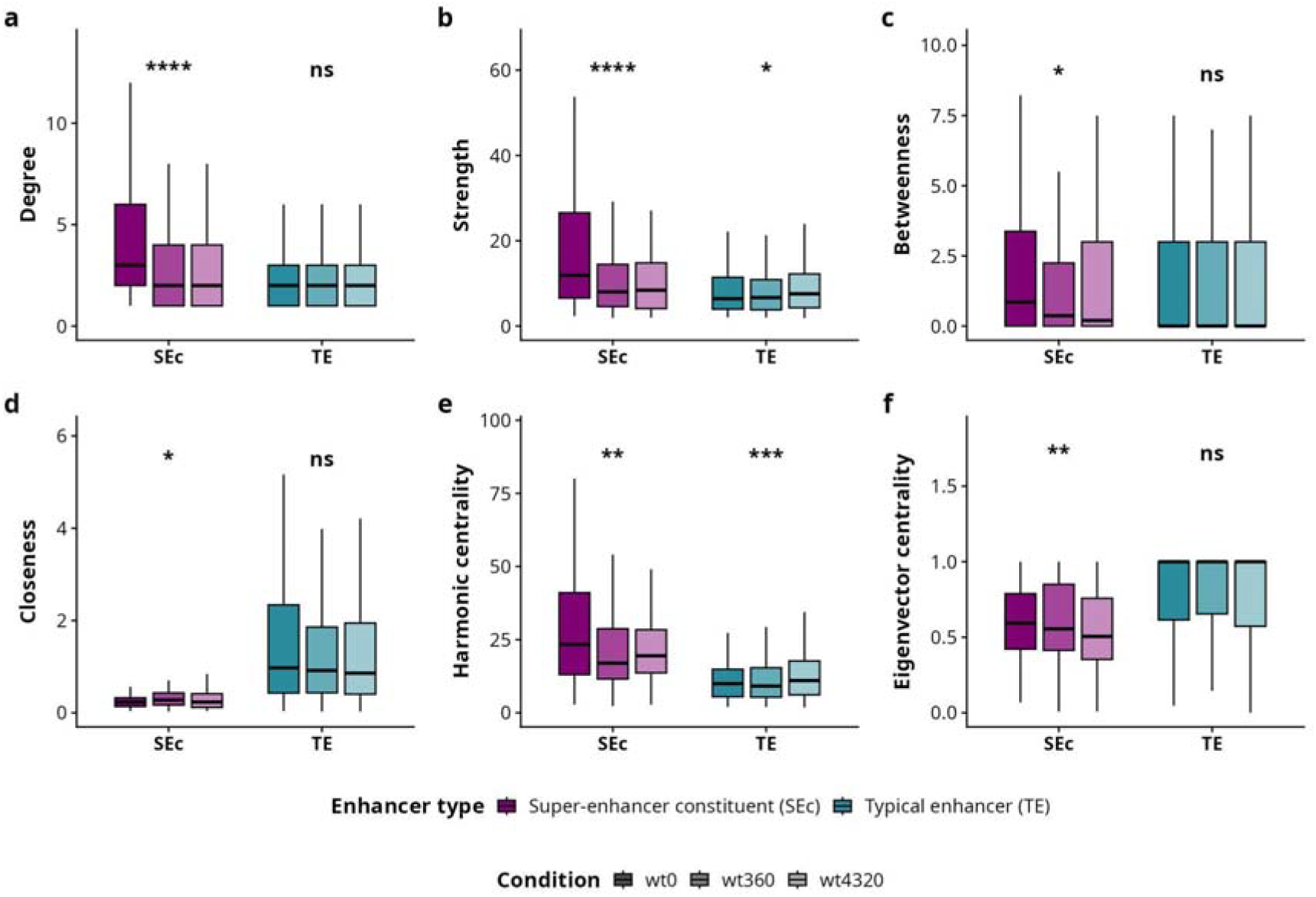
Distributions of SEc and TE (a) Degree, (b) Strength, (c) Betweenness, (d) Closeness, (e) Harmonic centrality, and (f) Eigenvector centrality over different conditions in SEc and TEs. Kruskal-Wallis test p<0.05 *, p<0.01 **, p<0.001 *** and p<0.0001 ****

To understand the main driving force behind the reorganization of the enhancer-promoter network during differentiation, analyses were performed to determine whether the change was driven by rearrangement of a persistent set of enhancers in the network (centrality change) or by turnover of enhancers themselves (enhancer change). For each pair of conditions, nodes were classified into shared (present in both conditions), gained (present only in the later condition), or lost (present only in the initial condition), and their proportions were computed relative to the union of nodes present in either condition, i.e., the total number of distinct nodes of that class appearing across the two conditions.

Although a substantial portion of both classes persisted between time points, approximately half of the nodes in each class were either gained or lost across each transition (Figure 3a; Supplementary Figure 7A, B for H9 and foreskin keratinocyte), suggesting that the enhancers are not merely rearranged; membership of the enhancers is substantially replaced during differentiation. This positions enhancer gain and loss as the principal force driving network change, consistent with a model in which differentiation reconfigures regulatory architecture largely by recruiting and retiring enhancers rather than by continuously rearranging the connections of a fixed set. (Stergachis *et al*. 2013)

**Figure 3:**
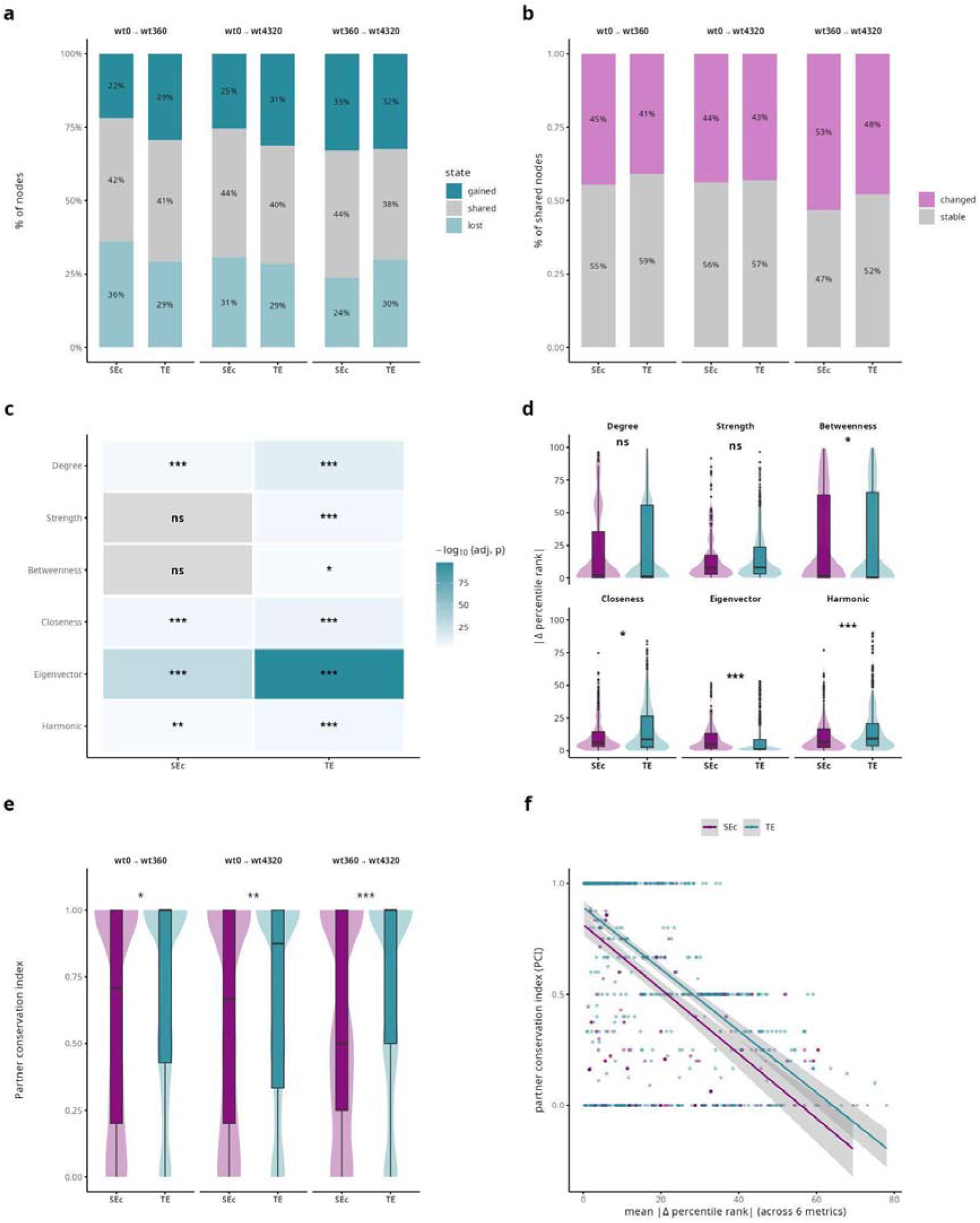
Representative figure showing (a) percentages of shared, gained, and lost SEc-anchored and TE-anchored loops between combinations of conditions; (b) percentages of SEc- and TE-anchored shared loops showing change in network properties over pairs of conditions; (c) heatmap showing the change in shared nodes across three conditions assessed using the Friedman test; (d) boxplot showing the magnitude of rank change of SEc and TE nodes between initial and final conditions; (e) Partner conservation index over combinations of conditions (f) correlation between the rank change and partner conservation index of SEc and TE. Comparisons assessed using Wilcoxon rank sum test p< 0.05 *, p<0.01 **, p<0.001 *** and p<0.0001 **** for (d) and (e)

We then examined the enhancers that persist across conditions to assess whether they retain their positions within the network. To make network properties comparable across networks of different sizes and densities, created from different conditions, we computed the six network centrality metrics (degree, strength, betweenness, closeness, eigenvector, and harmonic centrality) and converted each into a within-condition percentile rank, placing nodes on a common scale. Then, a node was considered to have changed if its rank shifted in at least 3 of the 6 metrics, and we found that a comparable proportion of SEc and TE nodes altered their network properties (Figure 3b). This was also observed in other differentiation systems assessed (Supplementary Figures 7C and 7D). To check for systematic movement in the ranks of network properties during the time course of differentiation, we first applied the Friedman test separately for SEc and TE for each metric (Figure 3c; Supplementary Figure 7 E, F). Both classes exhibited statistically significant systematic rank shifts on most metrics, confirming that the repositioning of nodes is a genuine phenomenon.

Second, to quantify the magnitude of repositioning, we computed the absolute change in percentile rank (|Δ rank|) between differentiation endpoints (wt0 and wt4320) and compared |Δ rank| for SEc with that for TE (Figure 3d). Median |Δ rank| for the majority of nodes and centrality metrics was low, suggesting that most persistent enhancers do not undergo major repositioning and that the position of enhancers within the network remains largely conserved; nevertheless, a proportion of enhancers were found to reposition. Most notably, eigenvector centrality showed a significant SEc–TE difference, indicating that the enhancers whose global network influence shifts most during differentiation are predominantly super-enhancers. This SEc–TE divergence in eigenvector centrality was reproduced in both supporting Hi-C datasets (h9, foreskin keratinocyte; Supplementary Figures 7g and 7h), indicating that it is a robust feature rather than a dataset-specific artifact. Therefore, it can be understood that many persistent nodes retain their relative network position, yet the shifts that do occur are systematic rather than random, indicating that a defined subset of retained enhancers undergo directed repositioning even as the majority hold steady.

Next, to assess partner identity directly, we calculated the partner conservation index (PCI) as the Jaccard similarity index between a node’s interaction partners across two conditions, and compared the PCI distributions between SEc and TE (Figure 3e). We found that across all three transitions, TE conserved its partners significantly better than SEc, and this difference intensified over the differentiation course (median PCI 0.71 vs 1.00 (wt0 -> wt360), *p* = 0.014; 0.50 vs 1.00, *p* = 1.1 × 10 (wt360->wt4320) for the successive transitions (SEc *vs.* TE)). Thus, despite comparable persistence of the two classes, the SEc that remain undergo more extensive partner rewiring than TE. This is compatible with the view of super-enhancers as dynamically regulated, cell-state-defining elements whose internal connectivity is continually reshaped, consistent with Hnisz et al. (2015), suggesting that SEc act as highly sensitive regulatory hubs during cell-state transitions (Hnisz *et al*. 2015).

Finally, we asked whether the two forms of change observed among persistent nodes, repositioning in the network and turnover of partners, are independent or reflect a common underlying event. Correlating each node’s overall centrality change (mean |Δ rank| across the six metrics) against its PCI revealed a strong negative relationship in both classes (Spearman ρ = −0.56 for SEc, ρ = −0.62 for TE; p < 10⁻30; Figure 3f). Persistent nodes that retained their partners also retained their network rank, while nodes that rewired their partners shifted rank in proportion. Partner turnover and centrality change are therefore not separable events but two readouts of the same remodeling process; when a persistent enhancer’s neighborhood turns over, its position within the network turns over with it.

This relationship is held in both supporting Hi-C datasets (Supplementary Table 2), indicating it is a general feature of differentiation-associated network remodeling rather than a K562-specific effect. (Supplementary Figure 7 i-l)

Alt text: Six-panel figure showing how the enhancer-promoter interaction network changes during PMA-induced differentiation. (a) Stacked bar plot showing the percentages of SEc and TE nodes that are shared, gained, or lost for each pair of conditions. About a quarter of the nodes are gained, a similar proportion are lost, and 40% are shared. (b) Stacked bar plots showing percentages of shared nodes that change or remain stable. Roughly 50% of the nodes changed, with a similar proportion for SEc and TE. (c) Heatmap of Friedman test results for six centrality metrics in SEc and TE, shaded by significance. Most metrics show significant changes. (d) Violin and box plots showing the absolute change in percentile rank between wt0 and wt4320 for each metric. Medians are low for both classes. SEc and TE differ significantly for betweenness, closeness, eigenvector, and harmonic centrality, but not for degree or strength. (e) Violin and box plots showing the partner conservation index for each condition pair. TE medians are near 1, while SEc medians are lower than 1. (f) Scatter plot of mean rank change against partner conservation index for each node. Both SEc and TE show a negative trend; nodes that lose their original partners shift in rank.

### 9.3 Functional consequences of enhancer promoter rewiring

Alt text: Four-panel figure showing pathways linked to enhancer-promoter loops that are gained or lost between untreated (wt0) and 72-hour PMA-treated (wt4320) K562 cells. (a) Venn diagram showing loops: 12,897 found only in wt4320, 10,317 only in wt0, and 1,261 shared. (b) Grouped bar chart showing enhancer-promoter loops by category. De novo gained: 67 SEc and 432 TE. Shared: 11 SEc and 68 TE. De novo lost: 65 SEc and 388 TE. (c) Dot plot of MSigDB Hallmark pathways enriched among promoters in each SEc and TE gained or lost category. Dot size indicates fold enrichment and color shows significance. (d) Horizontal bar chart of log2 fold enrichment, SEc over TE, for each pathway. Bars to the right indicate SEc bias: MYC targets V1 and V2, Wnt/β-catenin, E2F targets, and inflammatory response. Bars to the left are TE-specific: unfolded protein response, IL6/JAK/STAT3, glycolysis, complement, and adipogenesis.

We next wanted to investigate the functional impact of enhancer turnover; loops from the baseline and differentiated conditions were compared (Figure 4a) and categorized as gains or losses (Figure 4b). Each promoter set was tested for overrepresentation of MSigDB Hallmark programs (Liberzon *et al*. 2015) using the GREAT model (Gu and Hübschmann 2023), taking the genome as background.

**Figure 4:**
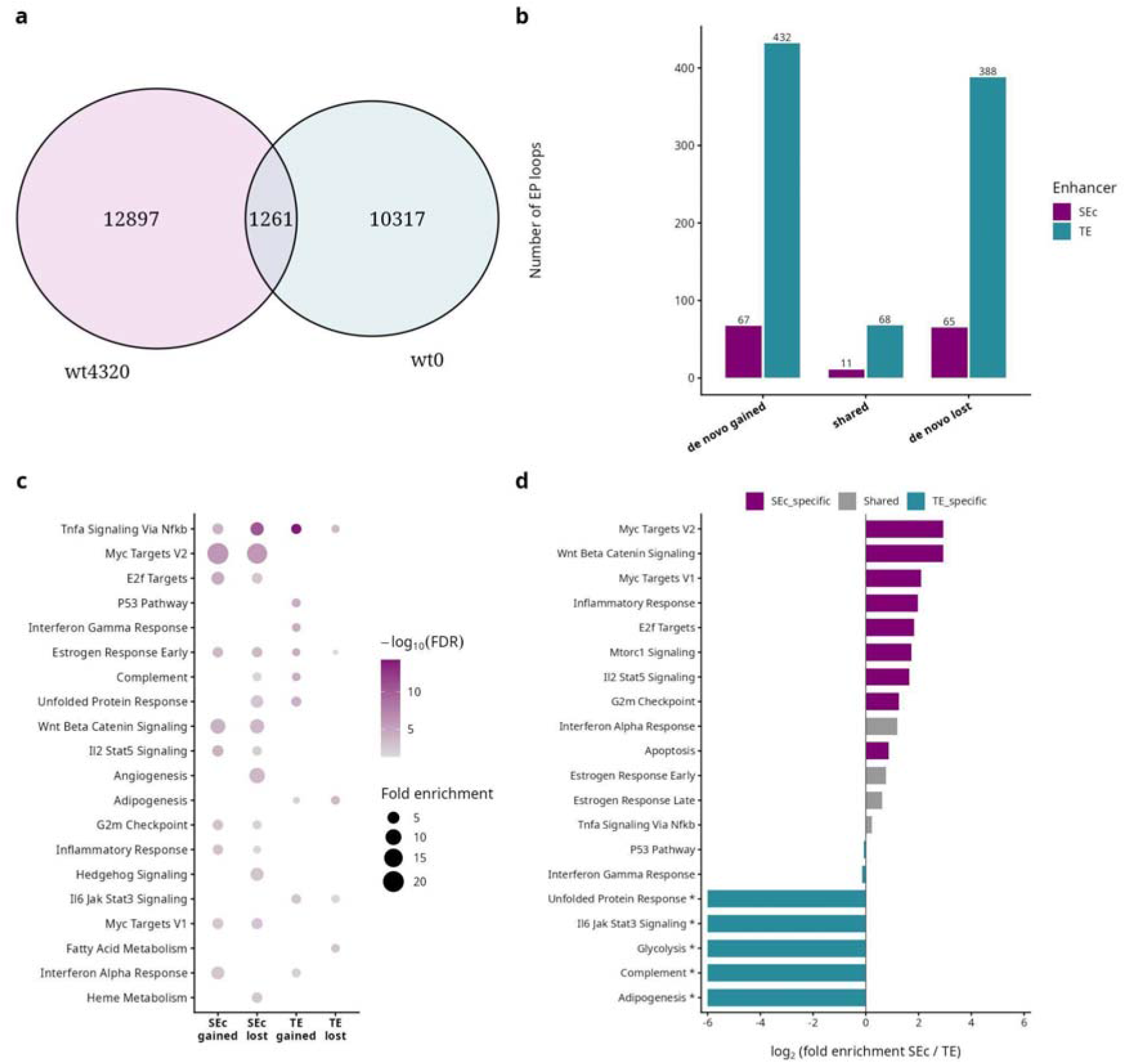
(a) Loops shared across baseline and final conditions (b) Distribution of SE-P loops and TE-P loops across the gain/loss categories (c) Pathways enriched for promoters controlled by SEc/TE across gain/loss categories (d) Bias of pathways enriched in SE-c vs TE.

Across all loop categories, TE-P loops substantially outnumbered SE-P loops, consistent with TEs forming the bulk of the enhancer-promoter network. The shared loops constituted only a small fraction of the total enhancer-promoter interactions, and given our earlier finding that network remodeling is driven predominantly by enhancer turnover, we focused the functional enrichment analysis on the de novo (gained and lost) loops, which capture the bulk of the regulatory reorganization. Across these de novo loops, promoters at SEc loops showed stronger pathway enrichment than those at TE loops (Figure 4c).

In the K562 cell line, promoters at de novo SEc loops were enriched for proliferation and growth-control programs, MYC targets (V1/V2), E2F targets, and Wnt/β-catenin signaling. On the other hand, promoters of de novo TE loops were enriched for stress- and immune-response programs, including interferon-γ response, complement, P53 pathway, and apoptosis (Figure 4c). Upon contrasting the enrichment of SEc vs TE pathways, the divergence was confirmed: MYC_TARGETS_V2, Wnt/β-catenin, and MYC_TARGETS_V1 were the most SEc-biased programs, whereas unfolded protein response, IL6/JAK/STAT3, glycolysis, complement, and adipogenesis were the most TE-biased (Figure 4d).

Together, this indicates a division of labor between enhancer classes in K562: SEc contacts are remodeled predominantly at proliferation and identity genes (MYC, E2F, cell-cycle targets), consistent with the established role of super-enhancers in driving high-output, cell-identity and proliferative programs (Hnisz *et al*. 2013, Lovén *et al*. 2013). Whereas TE loops turn over at metabolic and stress/immune loci such as glycolysis, adipogenesis, and complement.

The same analysis was also applied to two supporting datasets (h9 and foreskin keratinocyte) (Supplementary Figures 8 and 9). Two features were reproduced in both: TE-P loops numerically dominated every loop category (Supplementary Figures 8b and 9b), and SEc promoter sets again showed stronger enrichment than TE. However, the identity of SEc-associated programs was cell line specific rather than universal; in the h9 cell line, de novo gained SEc loops were enriched for metabolic programs (peroxisome, bile acid and fatty acid metabolism), and in foreskin keratinocytes for inflammatory/stress programs (TNFα-via-NFκB, hypoxia, estrogen response), rather than the MYC/E2F proliferative pathways seen in the K562 cell line.

This is consistent with super-enhancers being remodeled at the genes most central to each cell type’s identity and lineage, such that enhancer classes play an analogous structural role across systems while acting on distinct, context-appropriate gene programs.

## 10. Conclusion

In this study, we applied a network-based approach to represent Hi-C and HiChIP-derived loops as enhancer-promoter interaction networks and to analyze enhancer-promoter communication, distinguishing SEc from TE. In this study, the approach was used to characterize network organization and to analyze how it is remodeled during differentiation. Across multiple cell types and assays, the enhancer-promoter network was consistently modular, and clusters containing promoters, SEc, and TE were the largest, consistent with strongly regulated genes integrating multiple enhancer signals within discrete regulatory modules. Within this architecture, SEc and TE occupied distinct topological positions.

Applying this analysis to differentiation, we found that the network is reshaped primarily through enhancer turnover rather than through the rearrangement of a fixed set of enhancers. Among the enhancers that persisted, most retained their relative network position, but a defined subset underwent systematic repositioning, which was tightly coupled to turnover of interaction partners. SEc rewired their partners more extensively than TE did, consistent with SE acting as sensitive, cell-state-responsive elements. Linking the structural changes to function showed that SEc were focused on gene programs most central to each cell type’s identity, whereas TE contacts turned over more frequently at general metabolic and stress-response loci.

Overall, the recurrence of this organizational logic across cell types suggests it may represent a general principle of enhancer-promoter organization; beyond that, these findings were robust to annotation order, network projection, and choice of dataset, and similar analyses can be applied to study regulatory reorganization in other systems and cell-state transitions.

Some limitations and future directions of this work should be noted. Our analysis maps how enhancer-promoter contacts are wired and rewired, but it does not directly measure the transcriptional output of these changes; this would require integrating expression and perturbation data into the network to correlate topological changes with transcriptional changes. The network is also restricted to enhancer-promoter contacts and does not incorporate architectural elements or higher-order compartment structure, so it can capture only a single layer of chromatin communication rather than the full regulatory landscape. This can be addressed by extending this analysis to include architectural elements, higher-order features (TADs and compartments), and additional regulatory elements, allowing the enhancer-promoter network to be interpreted within its complete structural and regulatory context.

## Data Availability

The code underlying this article is available on GitHub (<u>github.com/ananya1607/SuperTypicalNetwork_analysis_code</u>)

## Author contributions statement

SAP performed data acquisition, contributed to the methodology, and prepared the initial draft of the manuscript. SR helped in conceptualization and formulation of the study. MK conceived the study, supervised the project, contributed to the methodology, and revised and finalized the manuscript.

## Acknowledgments

The authors thank the members of MKlab for the helpful discussions.

## Code availability and implementation

Code for implementation is available at https://github.com/ananya1607/SuperTypicalNetwork2.0

## Abbreviations

(SE): Super enhancer
(TE): Typical enhancer
(EP): enhancer-promoter
(SEc): Super enhancer constituent

## References

Belyaeva A, Venkatachalapathy S, Nagarajan M et al. Network analysis identifies chromosome intermingling regions as regulatory hotspots for transcription. Proc Natl Acad Sci 2017;114(52):13714–9. 10.1073/pnas.1708028115.

Bond ML, Davis ES, Quiroga IY et al. Chromatin loop dynamics during cellular differentiation are associated with changes to both anchor and internal regulatory features. Genome Res 2023;33(8):1258–68. 10.1101/gr.277397.122.

Csardi G. The Igraph Software Package for Complex Network Research. 2014. https://www.researchgate.net/publication/221995787.

Cunningham F, Allen JE, Allen J et al. Ensembl 2022. Nucleic Acids Res 2022;50(D1):D988–95. 10.1093/nar/gkab1049.

Grosveld F, Staalduinen J van, Stadhouders R. Transcriptional Regulation by (Super)Enhancers: From Discovery to Mechanisms. Annu Rev Genomics Hum Genet 2021;22(1):127–46. 10.1146/annurev-genom-122220-093818.

Gu Z, Hübschmann D. *rGREAT* : an R/bioconductor package for functional enrichment on genomic regions. Bioinformatics 2023;39(1):btac745. 10.1093/bioinformatics/btac745.

Harrison PW, Amode MR, Austine-Orimoloye O et al. Ensembl 2024. Nucleic Acids Res 2024;52(D1):D891–9. 10.1093/nar/gkad1049.

Hnisz D, Abraham BJ, Lee TI et al. Super-Enhancers in the Control of Cell Identity and Disease. Cell 2013;155(4):934–47. 10.1016/j.cell.2013.09.053.

Hnisz D, Schuijers J, Lin CY et al. Convergence of Developmental and Oncogenic Signaling Pathways at Transcriptional Super-Enhancers. Mol Cell 2015;58(2):362–70. 10.1016/j.molcel.2015.02.014.

Lawrence M, Huber W, Pagès H et al. Software for Computing and Annotating Genomic Ranges. PLoS Comput Biol 2013;9(8):e1003118. 10.1371/journal.pcbi.1003118.

Lee BH, Wu Z, Rhie SK. Characterizing chromatin interactions of regulatory elements and nucleosome positions, using Hi-C, Micro-C, and promoter capture Micro-C. Epigenetics Chromatin 2022;15(1). 10.1186/s13072-022-00473-4.

Liberzon A, Birger C, Thorvaldsdóttir H et al. The Molecular Signatures Database Hallmark Gene Set Collection. Cell Syst 2015;1(6):417–25. 10.1016/j.cels.2015.12.004.

Liu L, Ruan J. Utilizing networks for differential analysis of chromatin interactions. J Bioinform Comput Biol 2017;15(6). 10.1142/S021972001740008X.

Lovén J, Hoke HA, Lin CY et al. Selective inhibition of tumor oncogenes by disruption of super-enhancers. Cell 2013;153(2):320–34. 10.1016/j.cell.2013.03.036.

Lun ATL, Perry M, Ing-Simmons E. Infrastructure for genomic interactions: Bioconductor classes for Hi-C, ChIA-PET and related experiments. F1000Research 2016;5:950. 10.12688/f1000research.8759.2.

Malod-Dognin N, Pancaldi V, Valencia A et al. Chromatin network markers of leukemia. Bioinformatics 2020;36(Supplement_1):i455–63. 10.1093/bioinformatics/btaa445.

Mumbach MR, Rubin AJ, Flynn RA et al. HiChIP: efficient and sensitive analysis of protein-directed genome architecture. Nat Methods 2016;13(11):919–22. 10.1038/nmeth.3999.

Newman M. Networks, vol. 1. n.p.: Oxford University Press, 2018. 10.1093/oso/9780198805090.001.0001.

Oudelaar AM, Higgs DR. The relationship between genome structure and function. Nat Rev Genet 2021;22(3):154–68. 10.1038/s41576-020-00303-x.

Panigrahi A, O’Malley BW. Mechanisms of enhancer action: the known and the unknown. Genome Biol 2021;22(1):108. 10.1186/s13059-021-02322-1.

R Core Team. R: A Language and Environment for Statistical Computing. R Foundation for Statistical Computing, 24 June 2026.

Rao SSP, Huntley MH, Durand NC et al. A 3D Map of the Human Genome at Kilobase Resolution Reveals Principles of Chromatin Looping. Cell 2014;159(7):1665–80. 10.1016/j.cell.2014.11.021.

Rehemtulla A, Arndt A, Hart DA. Induction of plasminogen activator inhibitor type 2 expression during differentiation of human K562 cells towards a macrophage phenotype. Biochem Cell Biol 1990;68(12):1337–43. 10.1139/o90-196.

Reiff SB, Schroeder AJ, Kırlı K et al. The 4D Nucleome Data Portal as a resource for searching and visualizing curated nucleomics data. Nat Commun 2022;13(1):2365. 10.1038/s41467-022-29697-4.

Reyna J, Fetter K, Ignacio R et al. Loop Catalog: a comprehensive HiChIP database of human and mouse samples. Genome Biol 2025;26(1):175. 10.1186/s13059-025-03615-5.

Rowley MJ, Poulet A, Nichols MH et al. Analysis of Hi-C data using SIP effectively identifies loops in organisms from *C. elegans* to mammals. Genome Res 2020;30(3):447–58. 10.1101/gr.257832.119.

Schoenfelder S, Javierre BM, Furlan-Magaril M et al. Promoter Capture Hi-C: High-resolution, Genome-wide Profiling of Promoter Interactions. J Vis Exp June 2018;(136). 10.3791/57320.

Shlyueva D, Stampfel G, Stark A. Transcriptional enhancers: from properties to genome-wide predictions. Nat Rev Genet 2014;15(4):272–86. 10.1038/nrg3682.

Song S, Liu L, Feng C et al. SEdb 3.0: a comprehensive super-enhancer database across multiple species. Nucleic Acids Res 2026;54(D1):D322–33. 10.1093/nar/gkaf1294.

Stergachis AB, Neph S, Reynolds A et al. Developmental Fate and Cellular Maturity Encoded in Human Regulatory DNA Landscapes. Cell 2013;154(4):888–903. 10.1016/j.cell.2013.07.020.

Thandapani P. Super-enhancers in cancer. In: Pharmacology and Therapeutics. Elsevier Inc., July 2019, 199.129–38. 10.1016/j.pharmthera.2019.02.014.

Thibodeau A, Márquez EJ, Shin DG et al. Chromatin interaction networks revealed unique connectivity patterns of broad H3K4me3 domains and super enhancers in 3D chromatin. Sci Rep 2017;7(1). 10.1038/s41598-017-14389-7.

Uyehara CM, Apostolou E. 3D enhancer-promoter interactions and multi-connected hubs: Organizational principles and functional roles. Cell Rep 2023;42(4):112068. 10.1016/j.celrep.2023.112068.

Whyte WA, Orlando DA, Hnisz D et al. Master Transcription Factors and Mediator Establish Super-Enhancers at Key Cell Identity Genes. Cell 2013;153(2):307–19. 10.1016/j.cell.2013.03.035.

Zitnik M, Li MM, Wells A et al. Current and future directions in network biology. Bioinforma Adv 2024;4(1). 10.1093/bioadv/vbae099.

