## Supplementary Material for "Dynamic rewiring of super-enhancers shapes condition-specific genome regulation"

Supplementary Figures

**Supplementary Figure 1:**
Cluster sizes from K562 (untreated) bipartite network

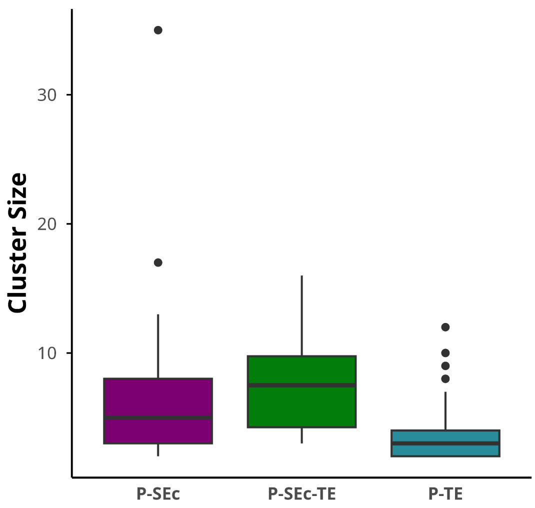

**Supplementary Figure 2:**

Cluster sizes across different cell lines (a) K562 (6hr post PMA treatment), (b) K562 (72hr post PMA treatment), (c) H9 cell line (untreated) (d) H9 cell line (day 2) (e) H9 cell line (day 5) (f) K562 cell line H3K27ac Hi-ChIP data (untreated) (g) MCF-7 cell line H3K27ac Hi-ChIP data (h) (i) (j)

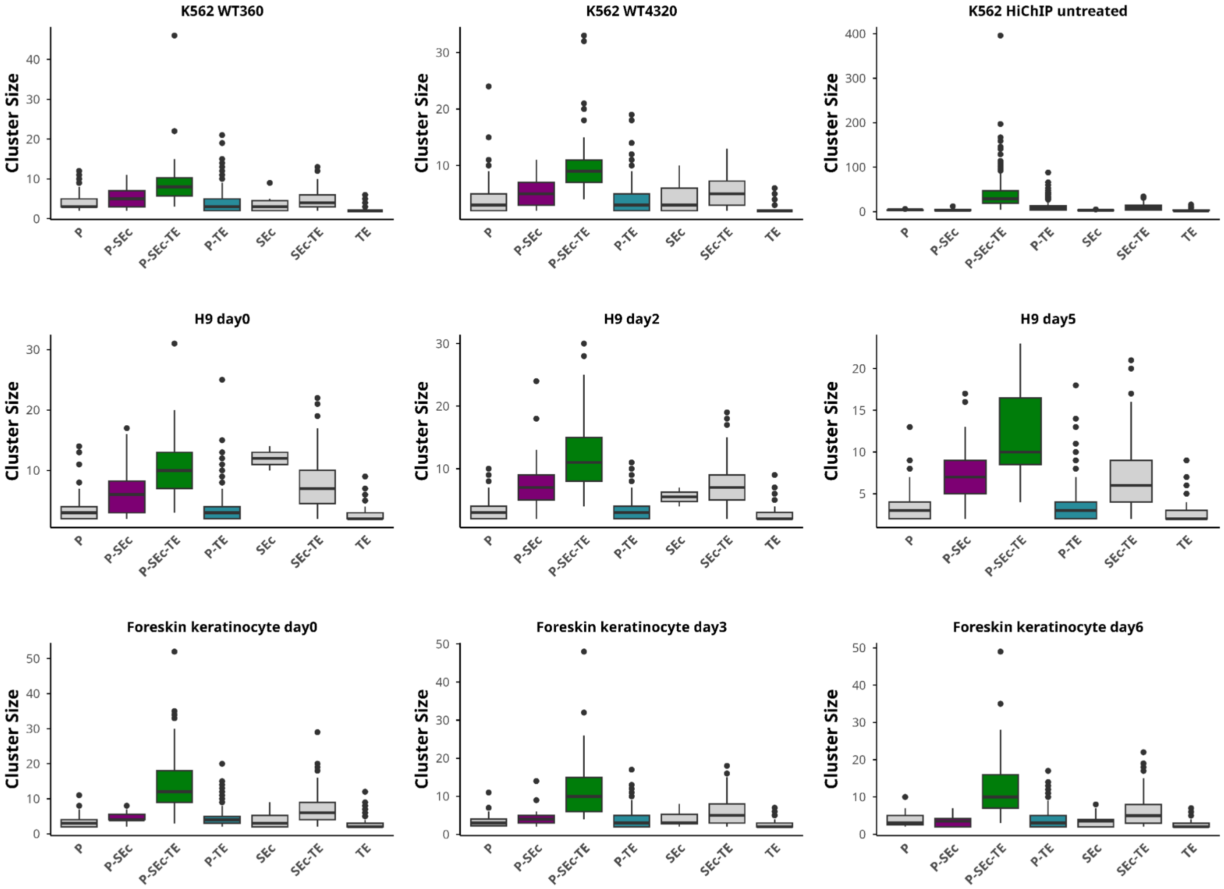

**Supplementary figure 3:**

Representative diagram showing the network-wide distributions of Degree , Strength, Betweenness, Closeness, Harmonic centrality, Eigenvector centrality between SE constituents and TEs. Wilcoxon rank sum test p<$0.05 *, p<0.01 ** , p<0.001 ***  and p<0.001 ***

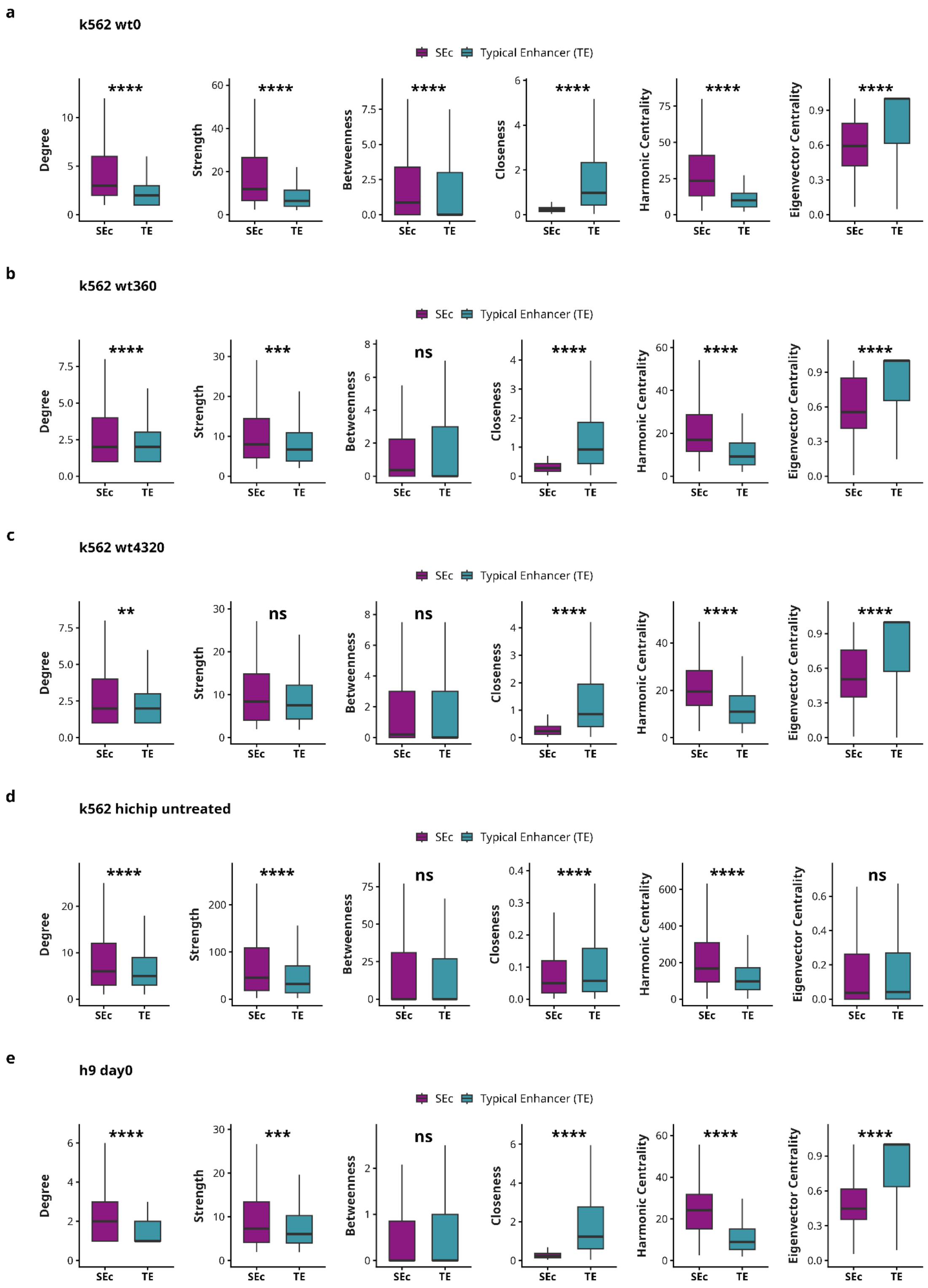

f

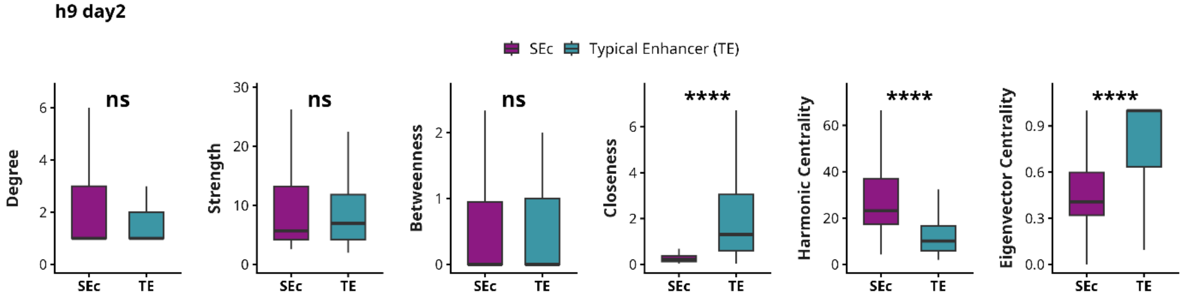
h9 day2
SEc
Typical Enhancer (TE)
Degree
Strength
Betweenness
Closeness
Harmonic Centrality
Eigenvector Centrality
ns
ns
ns
****
****
****
SEc
TE

g

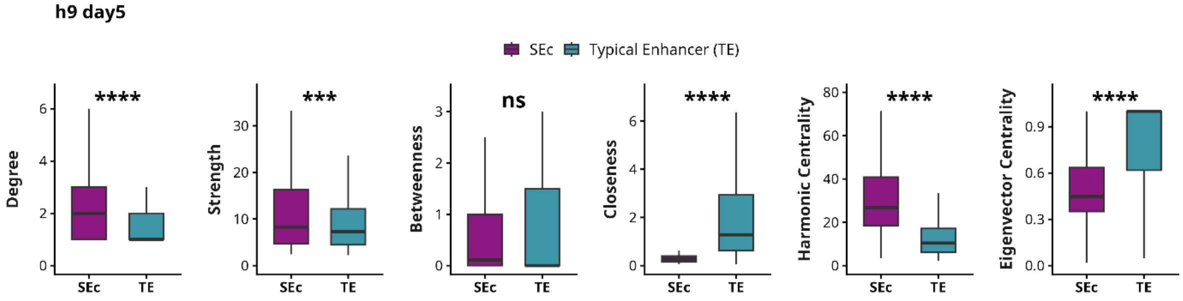
h9 day5
SEc
Typical Enhancer (TE)
Degree
Strength
Betweenness
Closeness
Harmonic Centrality
Eigenvector Centrality
****
***
ns
****
****
****
SEc
TE

h

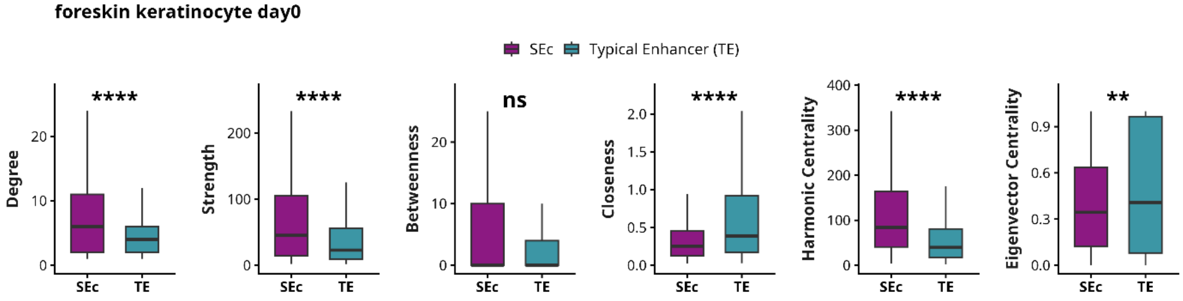
foreskin keratinocyte day0
SEc
Typical Enhancer (TE)
Degree
Strength
Betweenness
Closeness
Harmonic Centrality
Eigenvector Centrality
****
****
ns
****
****
**
SEc
TE

i

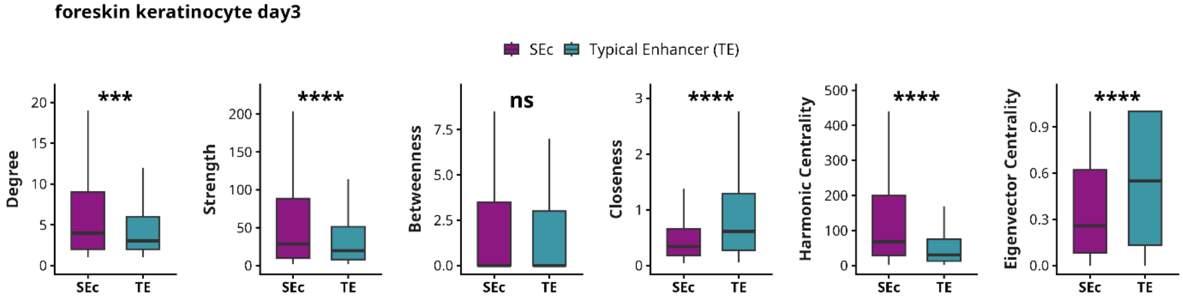
foreskin keratinocyte day3
SEc
Typical Enhancer (TE)
Degree
Strength
Betweenness
Closeness
Harmonic Centrality
Eigenvector Centrality
***
****
ns
****
****
****
SEc
TE

j

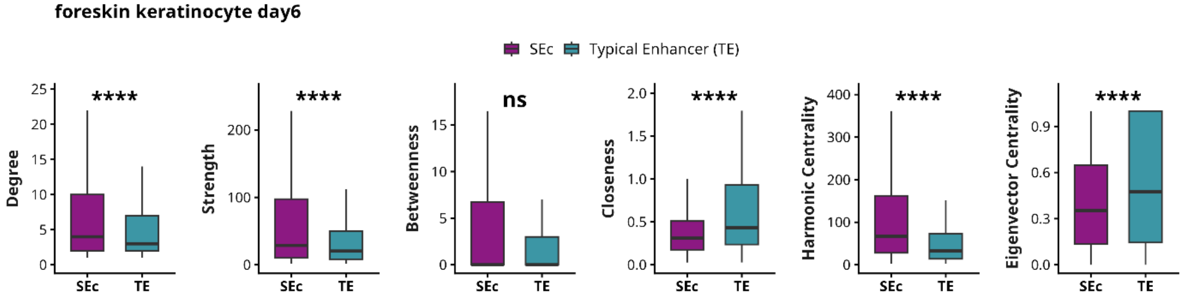
foreskin keratinocyte day6
SEc
Typical Enhancer (TE)
Degree
Strength
Betweenness
Closeness
Harmonic Centrality
Eigenvector Centrality
****
****
ns
****
****
****
SEc
TE

**Supplementary figure 4:**

Representative diagram showing the distribution of network properties in **bipartite networks** across cell lines (Wilcoxon rank sum test p<0.05 *, p<0.01 **, p<0.001 ***, p<0.0001 ****)

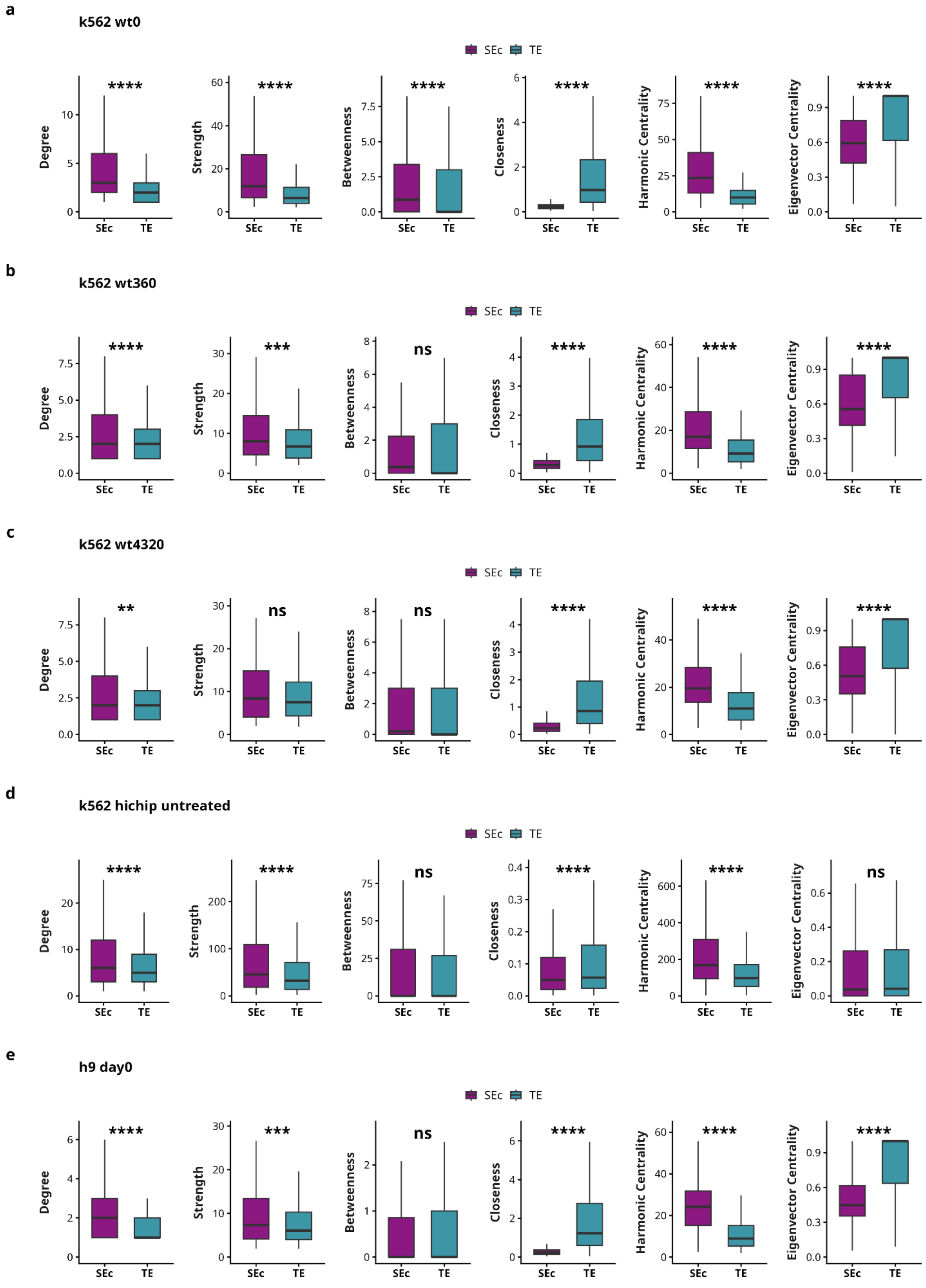

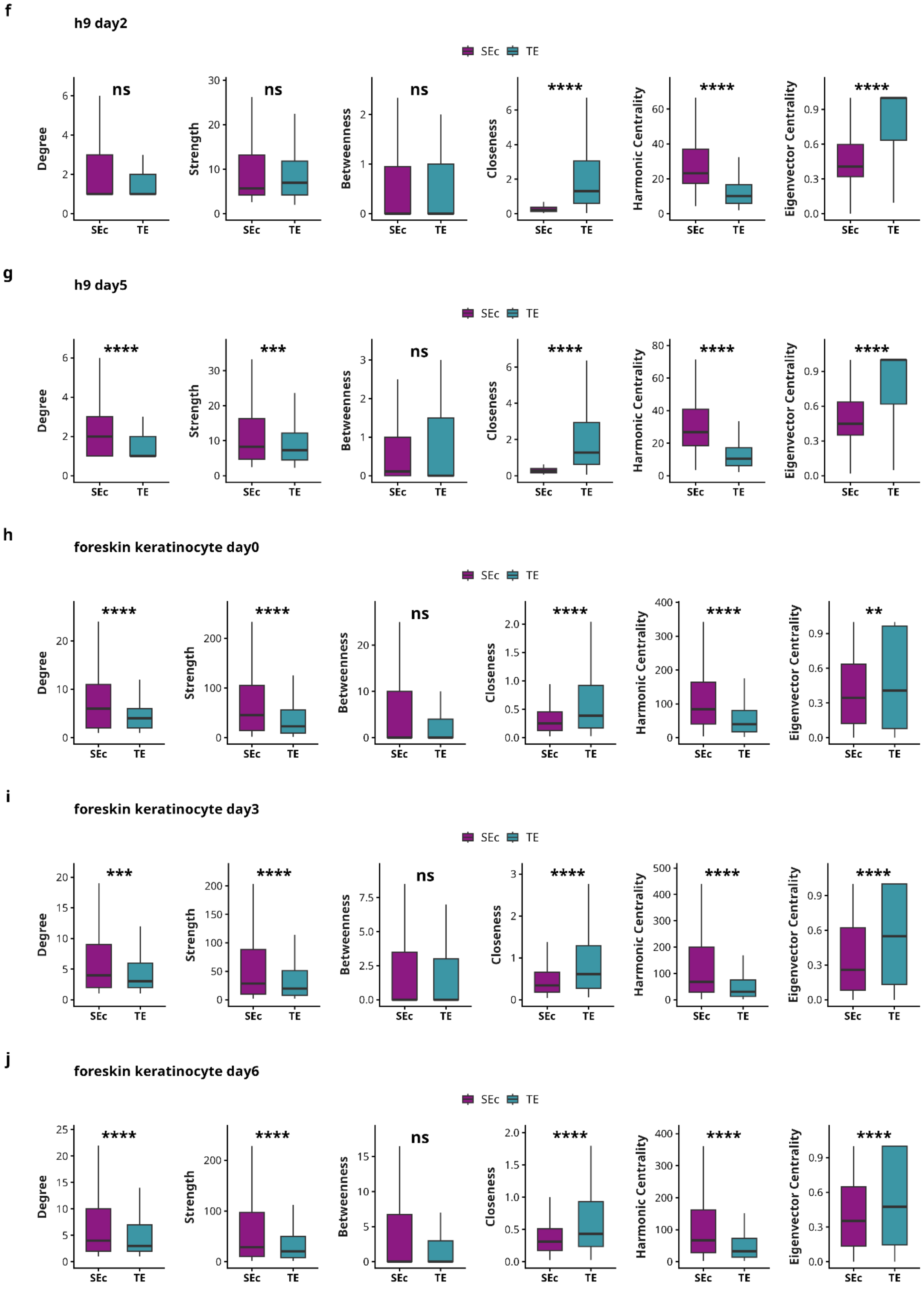

f
h9 day2
SEc TE
Degree ns
Strength ns
Betweenness ns
Closeness ****
Harmonic Centrality ****
Eigenvector Centrality ****
g
h9 day5
SEc TE
Degree ****
Strength ***
Betweenness ns
Closeness ****
Harmonic Centrality ****
Eigenvector Centrality ****
h
foreskin keratinocyte day0
SEc TE
Degree ****
Strength ****
Betweenness ns
Closeness ****
Harmonic Centrality ****
Eigenvector Centrality **
i
foreskin keratinocyte day3
SEc TE
Degree ***
Strength ****
Betweenness ns
Closeness ****
Harmonic Centrality ****
Eigenvector Centrality ****
j
foreskin keratinocyte day6
SEc TE
Degree ****
Strength ****
Betweenness ns
Closeness ****
Harmonic Centrality ****
Eigenvector Centrality ****

**Supplementary Figure 5:**
Representative diagram showing the distribution of network properties in networks created using **reverse priority** across cell lines (Wilcoxon rank sum test p<0.05 *, p<0.01 **, p<0.001 ***, p<0.0001 ****)

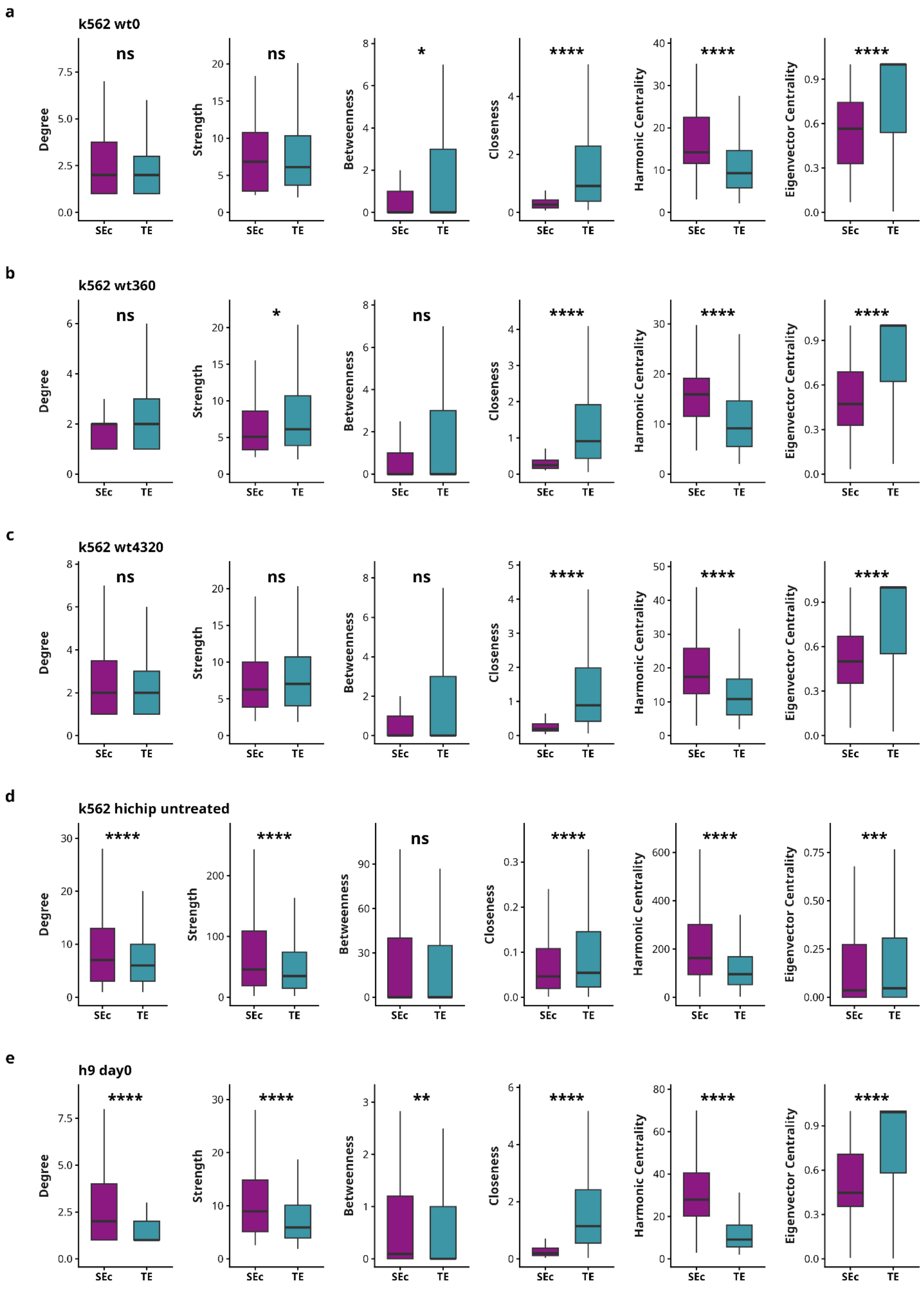

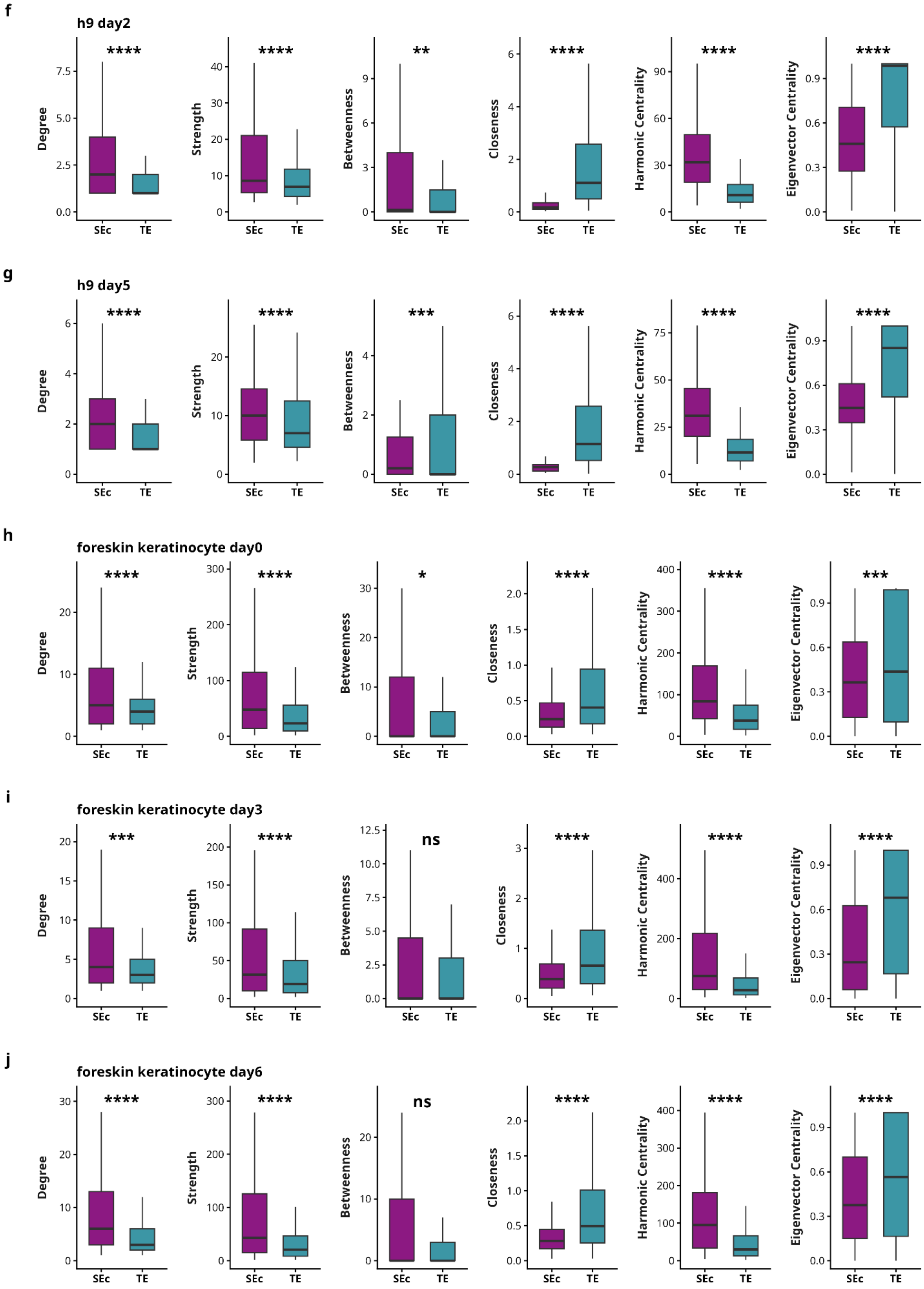
f
h9 day2
Degree
Strength
Betweenness
Closeness
Harmonic Centrality
Eigenvector Centrality
**** **** ** **** **** ****
SEc TE
g
h9 day5
Degree
Strength
Betweenness
Closeness
Harmonic Centrality
Eigenvector Centrality
**** **** *** **** **** ****
SEc TE
h
foreskin keratinocyte day0
Degree
Strength
Betweenness
Closeness
Harmonic Centrality
Eigenvector Centrality
**** **** * **** **** ***
SEc TE
i
foreskin keratinocyte day3
Degree
Strength
Betweenness
Closeness
Harmonic Centrality
Eigenvector Centrality
*** **** ns **** **** ****
SEc TE
j
foreskin keratinocyte day6
Degree
Strength
Betweenness
Closeness
Harmonic Centrality
Eigenvector Centrality
**** **** ns **** **** ****
SEc TE

**Supplementary Figure 6:**

(a-f) Distributions of SE constituent and TE (a) Degree, (b) Betweenness, (c) Closeness, (d) Eigenvector, and (e) Strength (f) Harmonic centrality over different conditions in super enhancer and typical enhancer in H9 cell line. (g-l) Distributions of SE constituent and TE (g) Degree, (h) Betweenness, (i) Closeness, (j) Eigenvector, and (k) Strength (l) Harmonic centrality over different conditions in super enhancer and typical enhancer in foreskin keratinocyte cell. Kruskal-Wallis test $p<0.05$ *, $p<0.01$ ** , $p<0.001$ *** and $p<0.0001$ ****

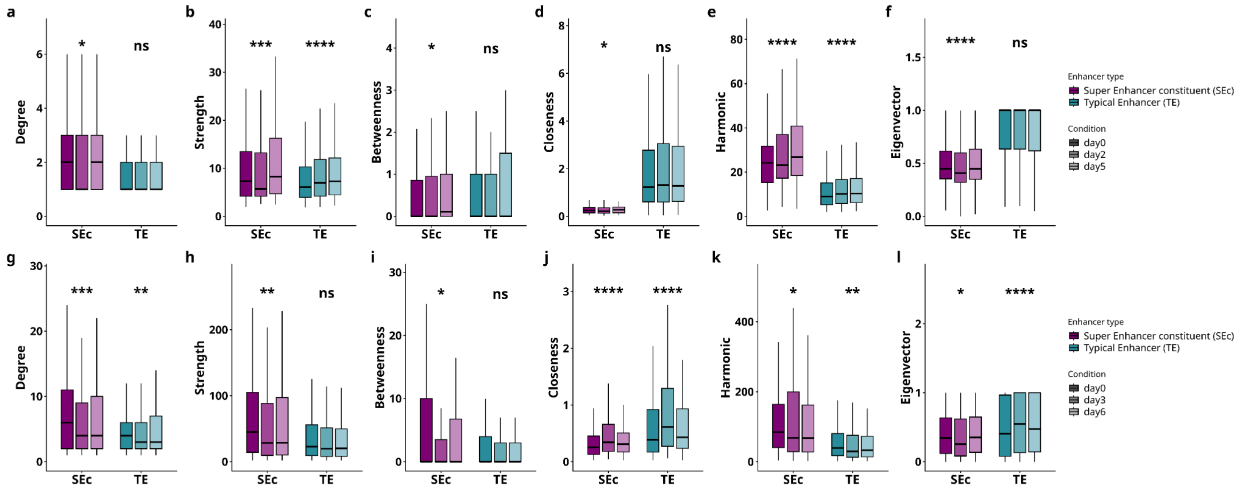

**Supplementary figure 7:**

Representative figure showing (a, b) percentages of shared, gained and lost SEc anchored and TE anchored loops between combinations of conditions in H9 cell line and foreskin keratinocyte cells respectively. (c, d) percentages of SEc and TE anchored shared loops showing change in network properties over pairs of conditions in H9 cell line and foreskin keratinocyte cells respectively. (e, f) heatmap showing the change of nodes shared nodes across three conditions assessed using friedman test in H9 cell line and foreskin keratinocyte cells respectively. (g, h) boxplot showing magnitude of rank change of SEc and TE nodes between initial and final conditions in H9 cell line and foreskin keratinocyte cells respectively. (i, j) Partner conservation index over combinations of conditions in H9 cell line and foreskin keratinocyte cells respectively. (k, l) correlation between the rank change and partner conservation index of SEc and TE comparison assessed using wilcoxon rank sum test in H9 cell line and foreskin keratinocyte cells respectively. (p< 0.05 *, p<0.01 ** , p<0.001 *** and p<0.0001 ****)

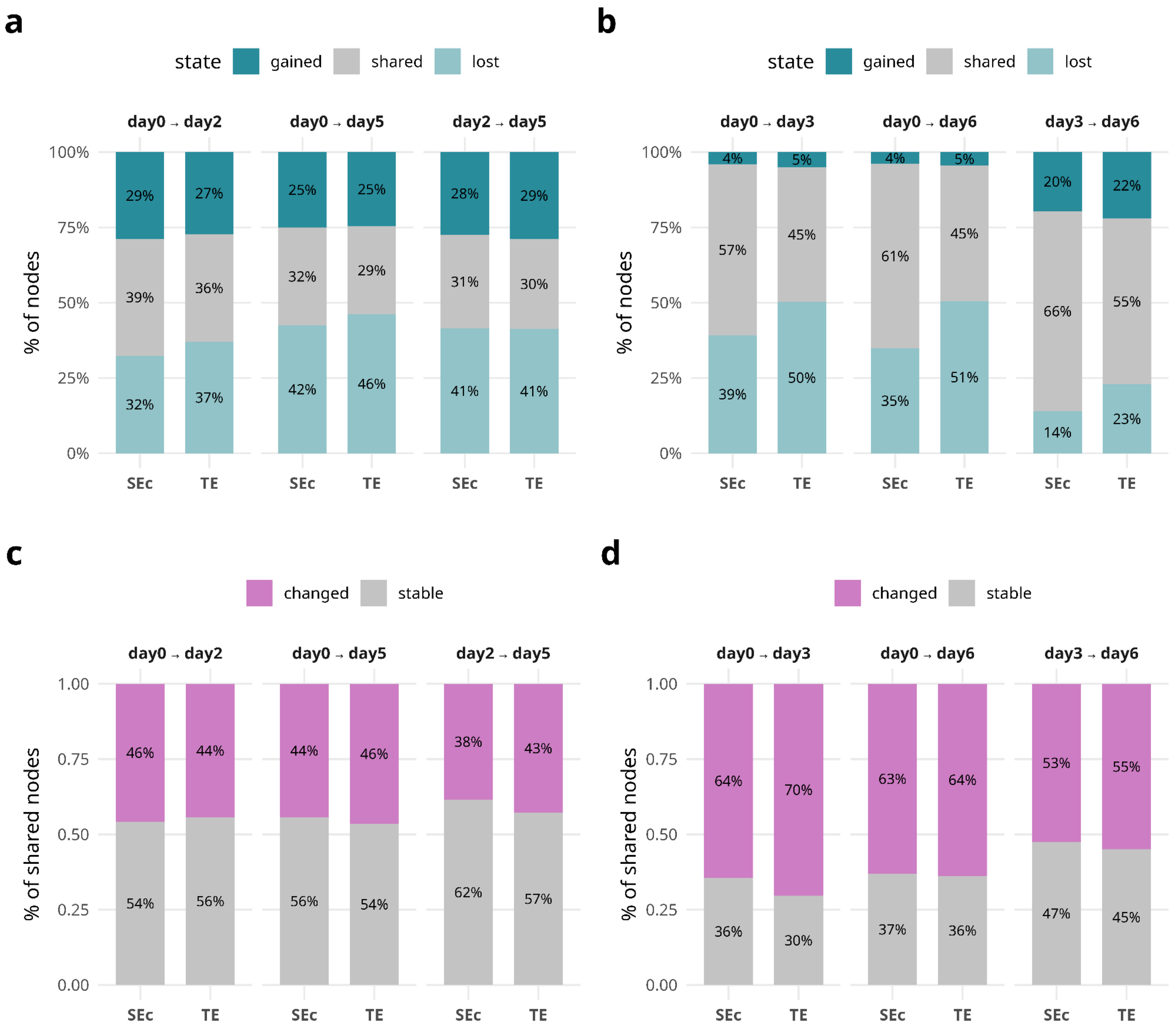

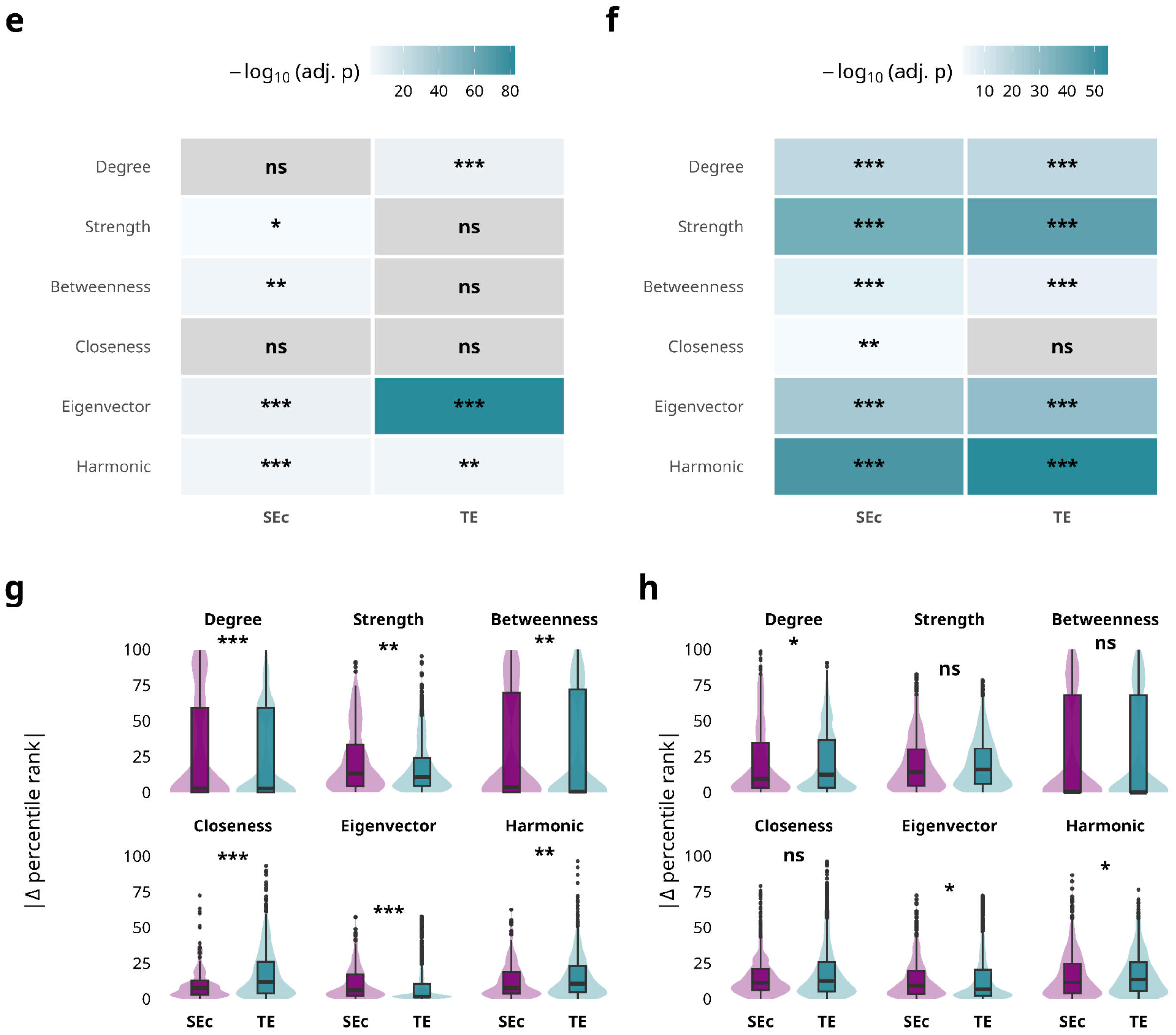

e
f
g
h
− log10 (adj. p)
20 40 60 80
10 20 30 40 50
Degree
Strength
Betweenness
Closeness
Eigenvector
Harmonic
SEc
TE
ns
***
*
**
|Δ percentile rank|
100
75
50
25
0

**i**

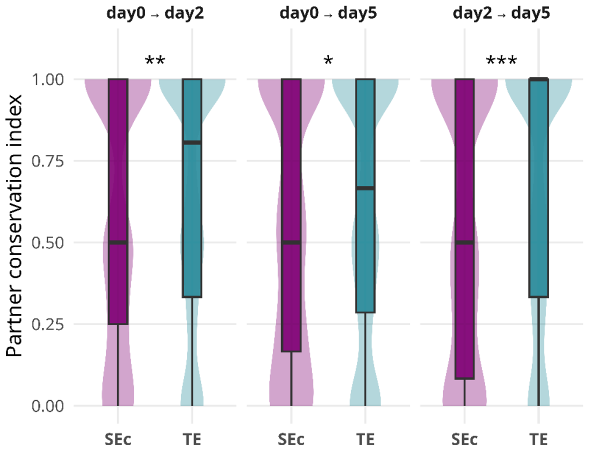
day0 → day2
day0 → day5
day2 → day5
**
*
***
Partner conservation index
1.00
0.75
0.50
0.25
0.00
SEc
TE
SEc
TE
SEc
TE

**j**

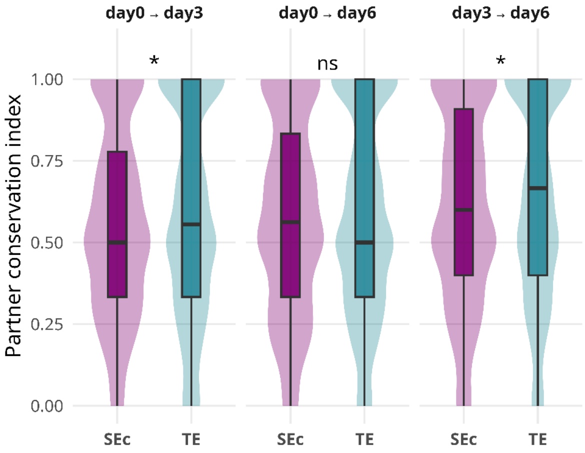
day0 → day3
day0 → day6
day3 → day6
*
ns
*
Partner conservation index
1.00
0.75
0.50
0.25
0.00
SEc
TE
SEc
TE
SEc
TE

**k**

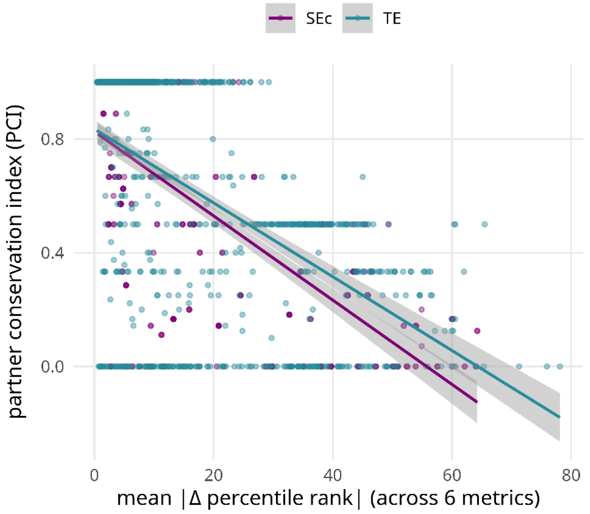
SEc
TE
partner conservation index (PCI)
0.8
0.4
0.0
0
20
40
60
80
mean |Δ percentile rank| (across 6 metrics)

**l**

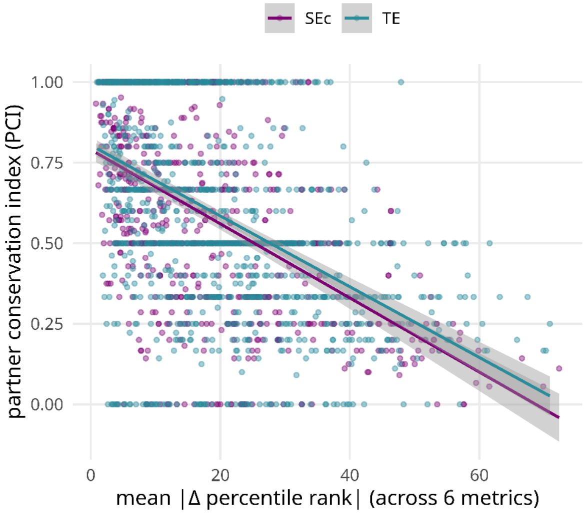
SEc
TE
partner conservation index (PCI)
1.00
0.75
0.50
0.25
0.00
0
20
40
60
mean |Δ percentile rank| (across 6 metrics)

**Supplementary figure 8:**

Representative diagram showing (A) Loops shared across baseline and final conditions (B) Distribution of SE-P loops and TE-P loops across the gain/loss categories (C) Pathways enriched for promoters controlled by promoters across gain/loss categories (D) Bias of pathways enriched in SE vs TE in H9 cells.

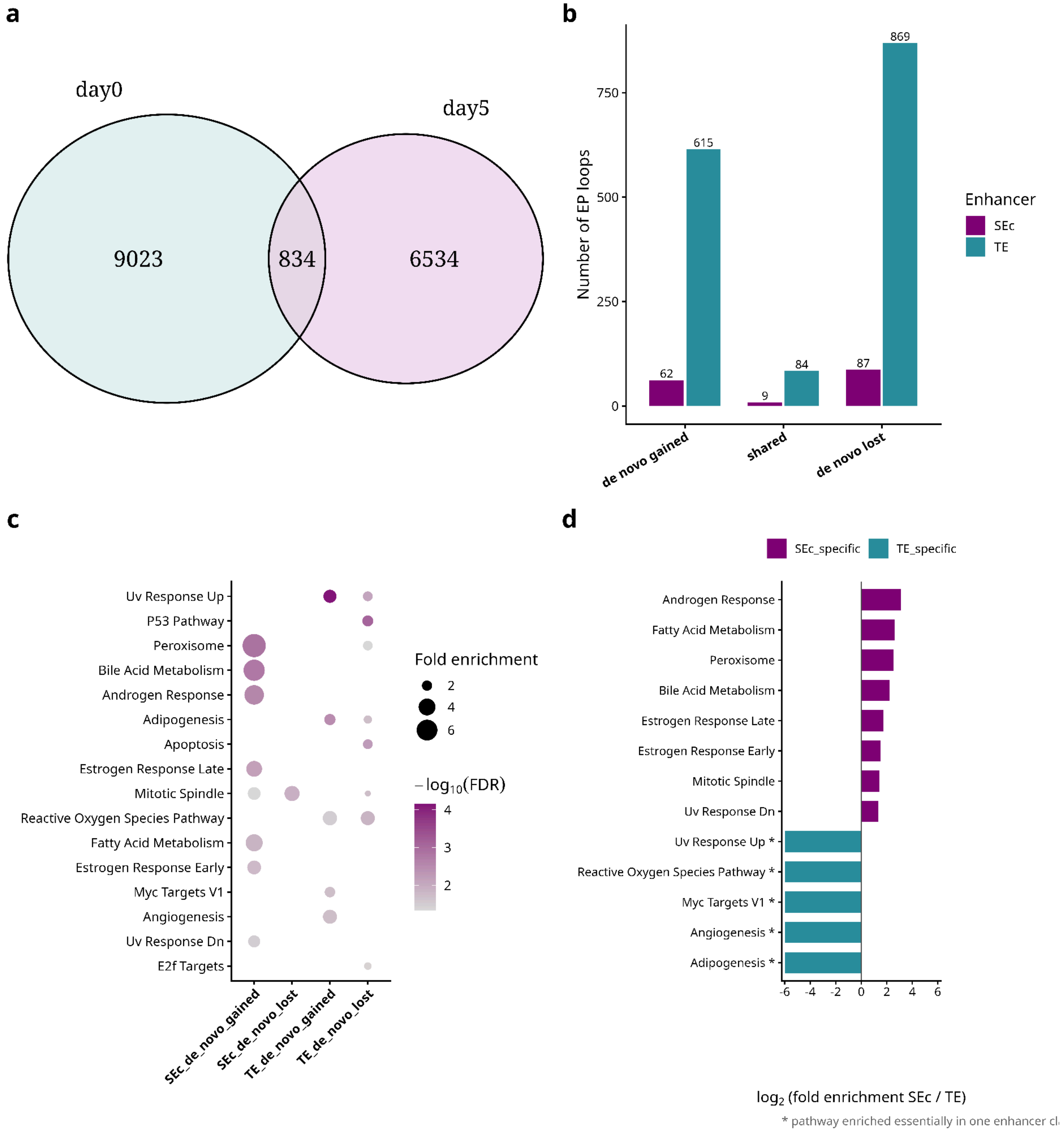

**Supplementary figure 9:**

Representative diagram showing (A) Loops shared across baseline and final conditions (B) Distribution of SE-P loops and TE-P loops across the gain/loss categories (C) Pathways enriched for promoters controlled by promoters across gain/loss categories (D) Bias of pathways enriched in SE vs TE in Foreskin Keratinocyte cells.

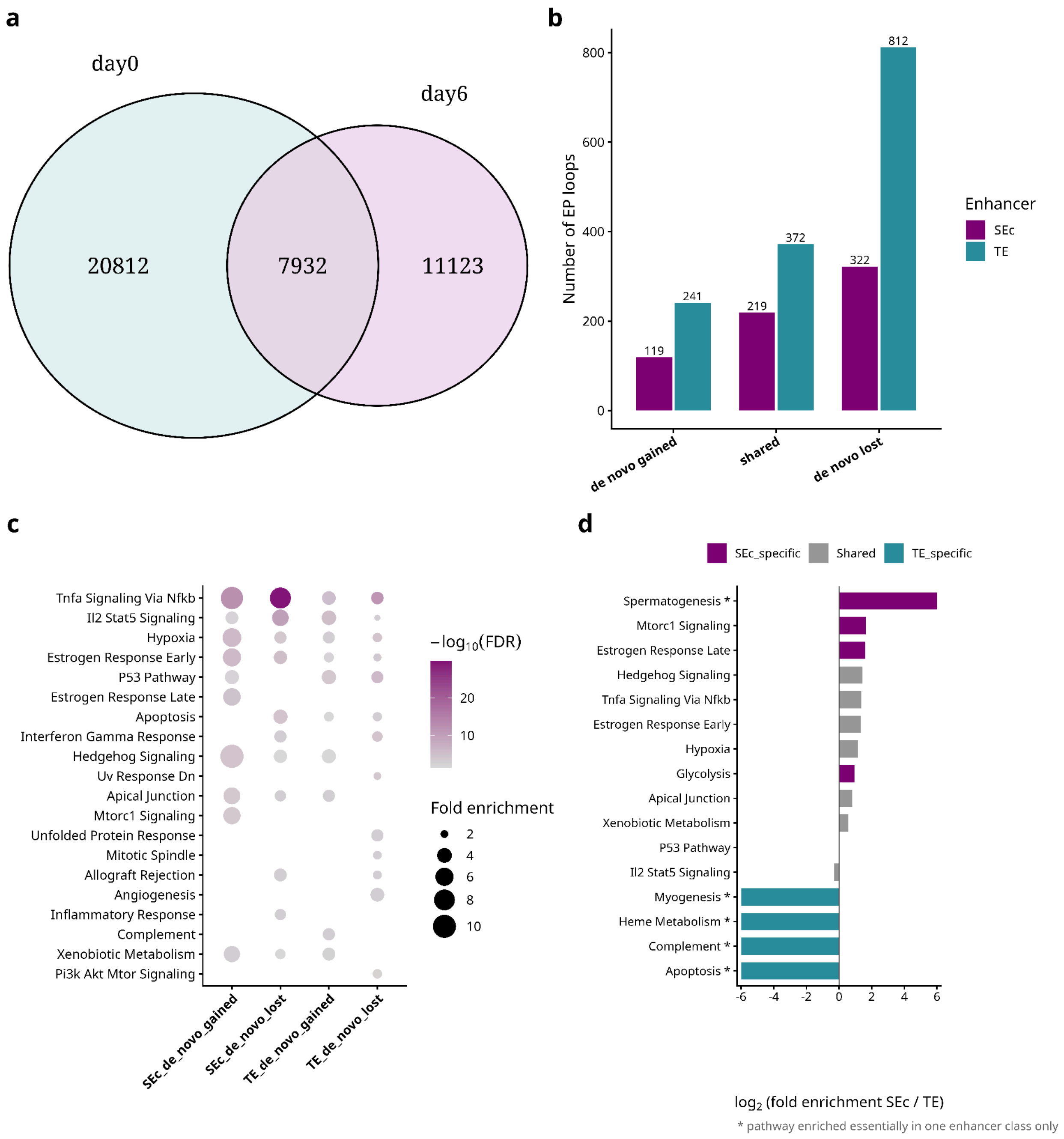
